# Evolutionary shaping of human brain dynamics

**DOI:** 10.1101/2022.06.07.495189

**Authors:** James C. Pang, James K. Rilling, James A. Roberts, Martijn P. van den Heuvel, Luca Cocchi

## Abstract

The human brain is distinct from those of other species in terms of size, organization, and connectivity. How do these structural evolutionary differences drive patterns of neural activity enabling brain function? Here, we combine brain imaging and biophysical modeling to show that the anatomical wiring of the human brain distinctly shapes neural dynamics. This shaping is characterized by a narrower distribution of dynamic ranges across brain regions compared with that of chimpanzees, our closest living primate relatives. We find that such a sharp dynamic range distribution supports faster integration between regions, particularly in transmodal systems. Conversely, a broad dynamic range distribution as seen in chimpanzees facilitates brain processes relying more on neural interactions within specialized local brain systems. These findings suggest that human brain dynamics have evolved to foster rapid associative processes in service of complex cognitive functions and behavior.

## INTRODUCTION

An important and unresolved problem in neuroscience is how connectivity, from neurons to macroscopic brain regions, can give rise to the complex dynamics that underlie behavior and advanced cognitive functions [1]. Identifying special features of the human brain that have evolved to support these complex neural dynamics is key in tackling this open question.

It is known that the human brain is approximately three times larger than would be expected in a primate with the same body mass [2,3]. Beyond general growth, neuroimaging analyses via magnetic resonance imaging (MRI) have indicated that a greater proportion of the human brain’s cortical surface is allocated to higher-order association cortices compared to primary sensory and motor areas [4–6]. This expansion of association areas is accompanied by increased anatomical connectivity [7], providing a structural substrate assumed to enable efficient region-to-region communication and integration of remote neural processes. Studies of the brain’s structural wiring, known as the *human connectome*, have shown widespread overlapping topological properties with those of other primates (like macaque and chimpanzees), accompanied by subtle but potentially consequential species differences [8].

Here, we ask how do the abovementioned structural changes shape whole-brain patterns of neural activity supporting brain function. To address this knowledge gap, we combine MRI data with advanced biophysical modeling to generate neural dynamics supported by the human connectome and the connectome of one of our closest living primate relatives: the chimpanzee. The use of validated biophysical models is crucial to tease apart and explain the neural basis of inter-species differences in whole-brain function, which cannot be achieved with current neuroimaging techniques [9]. By combining this innovative approach with a unique cross-species dataset, we reveal core neural principles likely to explain differences in brain function between humans and non-human primates.

## RESULTS

### Modeling neural dynamics

We begin by creating the connectomes of humans and chimpanzees. We use unique diffusion MRI data for adult humans (*Homo sapiens*) and sex-matched and age-equivalent chimpanzees (*Pan troglodytes*) to reconstruct the connectomes [7,10]. The connectomes represent cortico-cortical structural connections between 114 species-matched regions in both hemispheres (Table S1) from which we create group-averaged weighted human and chimpanzee connectomes [10] (Fig. 1A). Next, we combine the connectomic data with a validated biophysical (generative) model [11–14] (Fig. 1B; see STAR Methods) to generate regional neural activity across time (*neural dynamics*) specific to each species. This model has been shown to reproduce empirical human functional neuroimaging data [13,14], which we confirmed (Fig. S1). Notably, we also confirmed the model’s suitability to match non-human primate data (Fig. S1). These validations of the model on human and non-human primate data are important to ensure that the outcomes of the model capture meaningful properties of brain activity.

**Fig. 1.**
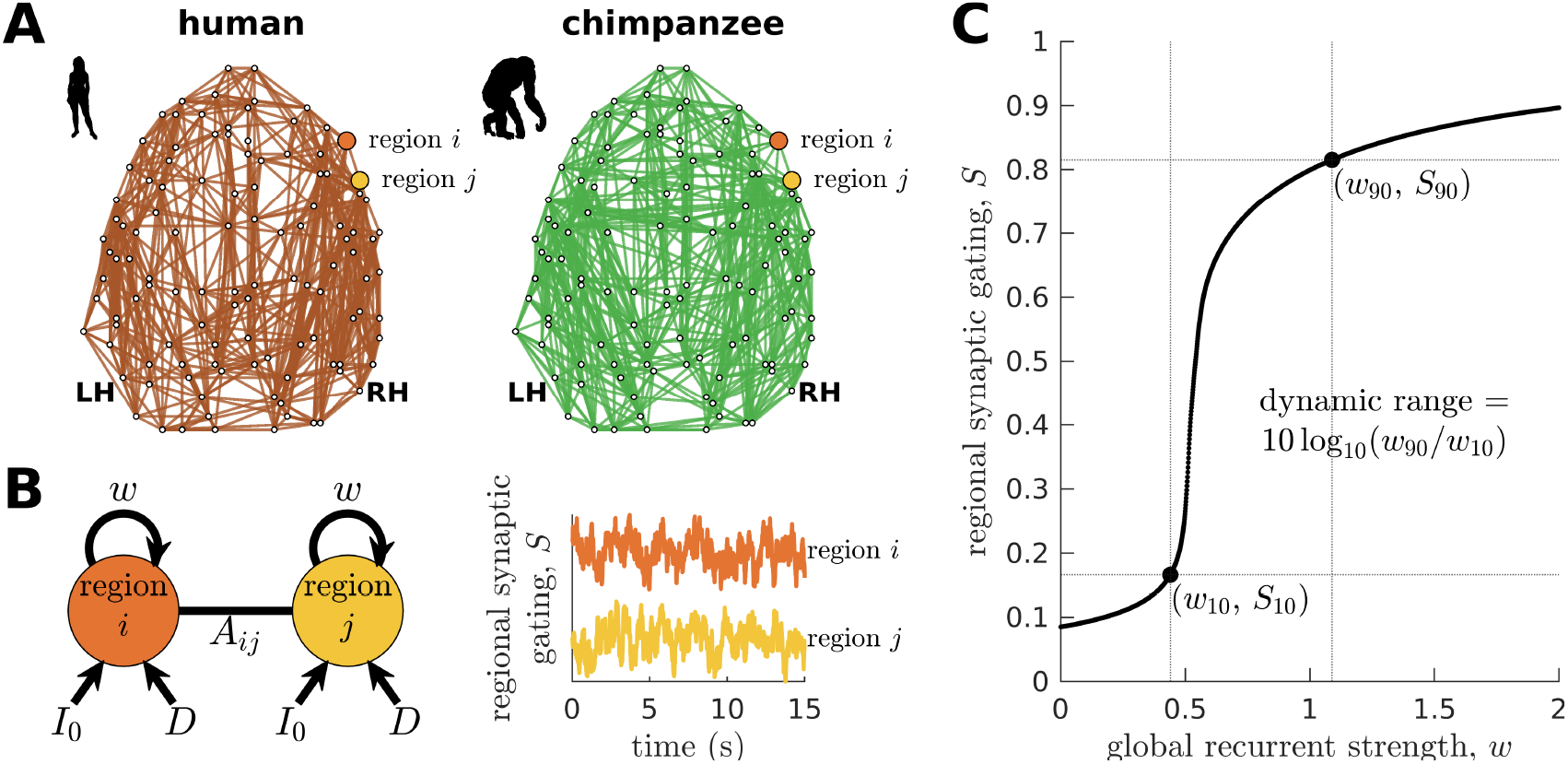
Human and chimpanzee connectomes and brain network modeling. (**A**) Group-averaged connectomes of humans and chimpanzees visualized on the same brain template. Top 20% of connections by strength are shown. (**B**) Schematic diagram of the model. Each brain region is recurrently connected with strength *w* and driven by an excitatory input *I*_0_ and white noise with standard deviation *D*. The connection between regions *i* and *j* is weighted by *A_ij_* based on the connectomic data. The regional neural dynamics are represented by the synaptic gating variable *S*; high S translates to high neural activity. (**C**) Method for calculating the dynamic range of each brain region. Note that *S_x_* = *S*_min_ + (*x*/100) * (*S*max – *S*_min_), with *w_x_* being the corresponding global recurrent strength at *S_x_* and *x* = {10,90}.

To understand how whole-brain activity patterns emerge, we analyze the intrinsic characteristics of regional neural dynamics. In particular, we determine a brain region’s response function, describing the up- or downregulation of its activity following global (brain-wide) modulations in the strength of recurrent connections (Fig. 1C). This process can be linked to how structures in the ascending neuromodulatory systems (e.g., noradrenergic) facilitate the reorganization of cortex-wide dynamics by allowing coordinated communication between otherwise segregated systems [15–17]. In particular, neuromodulatory agents can mediate changes in the excitability of cortical regions, which can occur in a subsecond timescale [18], effectively driving modulations in the strength of recurrent connections in cortical regions (i.e., our model’s *w* parameter). We measure the extent of this regulation in terms of the *neural dynamic range*, such that high (diffuse) dynamic range means that a region can respond to a wide range of changes in excitability. Conversely, a low (sharp) dynamic range means that a region can quickly transition between low and high levels of activity with even small changes in excitability. Note that our dynamic range is based on an excitability-output function (Fig. 1C) rather than an input stimulus-output function commonly used in previous studies [19]. Brain regions with similar dynamic ranges are more likely to coactivate, allowing efficient region-to-region integration of neural processes. Using this property, we aim to reveal key principles of whole-brain neural dynamics setting humans apart from other species.

### Humans have more constrained neural dynamics than chimpanzees

We find that the response functions of human brain regions (reflecting how activity changes versus modulations in global excitability) are more similar to one another compared to those of chimpanzees (Fig. 2A). We quantitatively test this observation by calculating the distribution of dynamic ranges across regions (Fig. 2B). The results show that the human brain has neural dynamic ranges characterized by a narrower distribution (standard deviation *σ* = 0.12) as compared to the chimpanzee brain (*σ* = 0.48). This finding is robust against differences in individual-specific connectomes (Figs. S2A and S2B), brain volume (Fig. S2C), connection density (Fig. S3), inter-individual variability of connection strengths (Fig. S4), data sample size (Fig. S5), propagation delays between brain regions (Fig. S6), and heterogeneous excitatory input across brain regions (Fig. S7). Moreover, our results are replicated on independent human data from the Human Connectome Project [20] (Fig. S8) and a different computational model (Fig. S9).

**Fig. 2.**
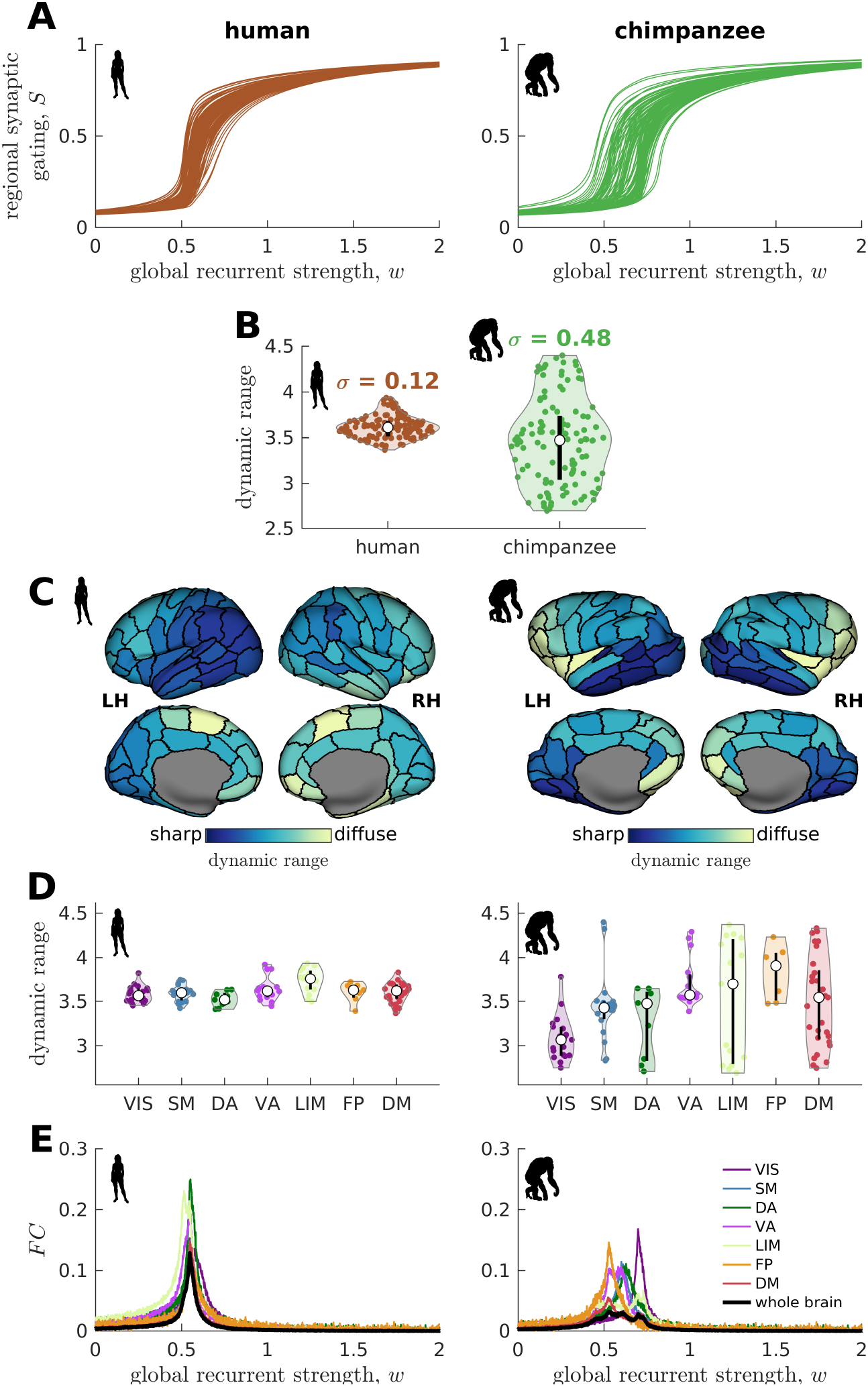
Human and chimpanzee neural dynamics. (**A**) Regional neural dynamics as a function of global recurrent strength (*w*). (**B**) Violin plot of the distribution of dynamic ranges across brain regions. Each violin shows the first to third quartile range (black line), median (white circle), raw data (dots), and kernel density estimate (outline). *σ* is the standard deviation of the distribution. (**C**) Spatial organization of dynamic ranges. Data are visualized on inflated cortical surfaces. Light color represents diffuse (high) dynamic range and dark color represents sharp (low) dynamic range. (**D**) Violin plot of the distribution of dynamic ranges in 7 canonical brain networks. Violin plot details are similar to those in panel B. (**E**) Simulated average functional connectivity (FC) within the networks in panel D as a function of *w*. The black line represents the average *FC* across the whole brain.

### Neural dynamic range is spatially organized along the anterior-posterior brain axis

When we map the dynamic ranges onto the anatomical locations of each brain region, we find that both species follow a dominant gradient of neural dynamic ranges spatially organized along the anterior-posterior axis (Fig. 2C). Specifically, anterior regions (e.g., frontal regions known to be expanded in humans compared to chimpanzees [3,4]) show neural dynamics with higher (more diffuse) dynamic ranges, while posterior regions (e.g., occipital cortex known to be relatively similar in size across the two species) have lower (sharper) dynamic ranges. Interestingly, we observe that this dominant gradient is more prominent in chimpanzees than in humans (Fig. S10).

### Similar neural dynamic ranges across regions enables brain network integration

We next ask whether brain regions belonging to specific functional networks have neural dynamics with similar levels of dynamic range. We cluster brain regions into 7 common large-scale brain networks according to [21,22]: Visual (VIS), Somatomotor (SM), Dorsal-Attention (DA), Ventral-Attention (VA, also known as the Salience network), Limbic (LIM), Frontoparietal (FP), and Default-Mode (DM) networks (Fig. S11). These networks represent functionally coupled regions across the cerebral cortex. In humans, brain regions belonging to each functional network show relatively similar neural dynamic ranges (Fig. 2D). Conversely, in chimpanzees, neural dynamic ranges follow a marked functional hierarchy with cognitive networks (i.e., VA, LIM, FP, DM) having higher median values than sensory networks (i.e., VIS, SM, DA). Furthermore, the patterns of within-network changes in functional connectivity versus modulations in global excitability overlap strongly in humans but not in chimpanzees (Fig. 2E). Thus, similar levels of regional dynamic ranges allow the human brain to better integrate activity within functionally specialized brain networks (colored lines in Fig. 2E) and the whole brain (black line in Fig. 2E). This finding is consistent with the higher level of structural integration imposed by the human connectome, as quantified by graph theoretical measures of path length (Fig. S12).

### Neural dynamic range differentiates humans and non-human primates

To further test the hypothesis that neural dynamic range is a key feature setting the human brain apart from the brains of other species, we perform similar analyses on other non-human primate connectomic data: macaque (*Macaca mulatta*) [23] and marmoset (*Callithrix jacchus*) [24]. Neural dynamics are obtained via the model in Fig. 1B and using weighted connectomes generated from diffusion MRI (for macaques) and invasive tract tracing (for marmosets). The connectomes represent connections between 82 and 55 regions of the macaque and marmoset brains, respectively. Both species have neural response functions and dynamic ranges closer to chimpanzees than to humans (Fig. 3; *σ* = 0.23 for macaque and *σ* = 0.31 for marmoset), further validating our results in Fig. 2 across species and across methodological differences in connectome data type and resolution. To verify that the macaque results are not driven by one apparent outlier, as seen in Fig. 3A, we perform the analysis on independent macaque dataset and find that the results are replicated (Fig. S13).

**Fig. 3.**
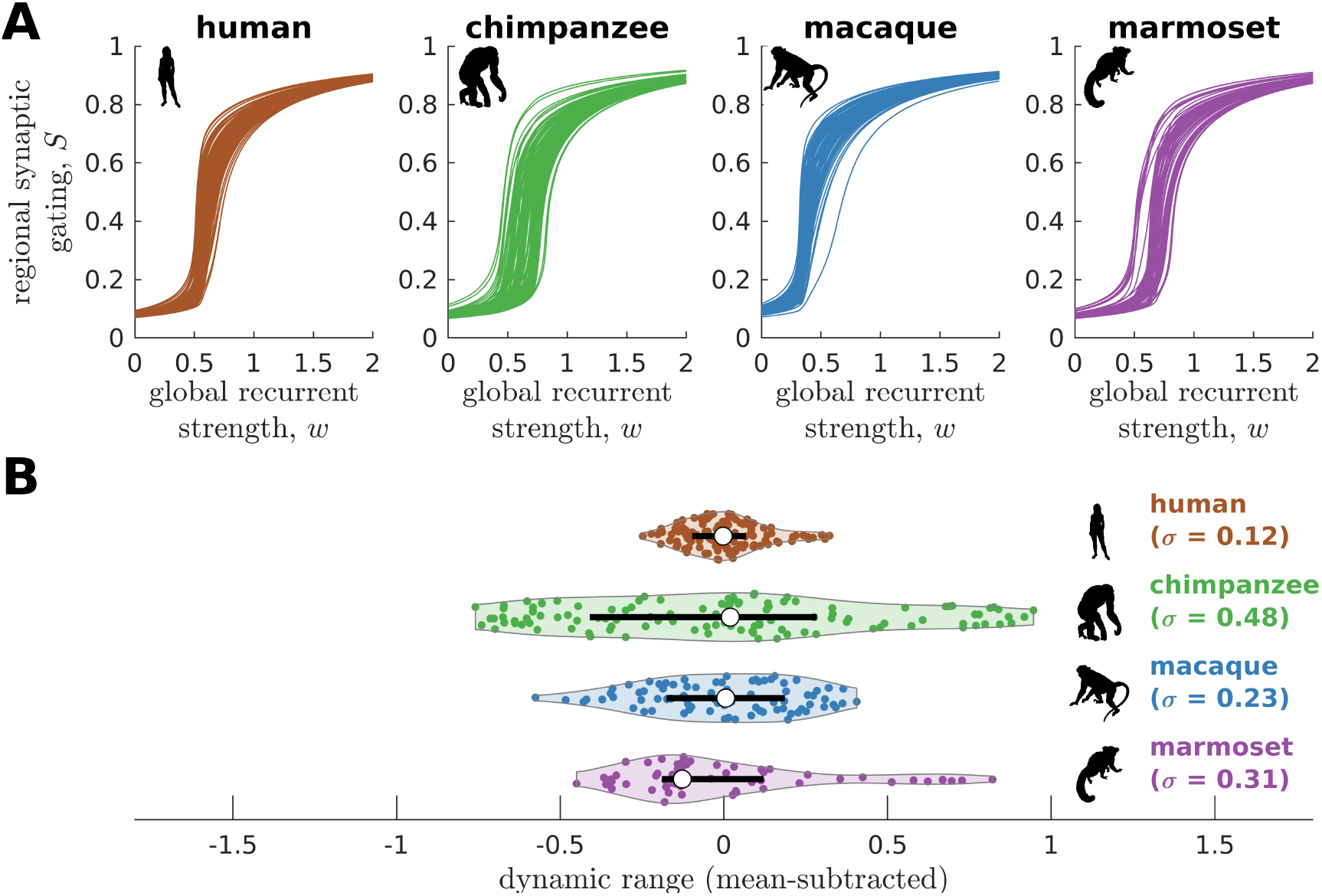
Neural dynamics of human and non-human primates. (**A**) Regional neural dynamics as a function of global recurrent strength (*w*) for human, chimpanzee, macaque, and marmoset. (**B**) Violin plot of the distribution of dynamic ranges across brain regions. Violin plot details are similar to those in Fig. 2B. The data are mean-subtracted for visual purposes. *σ* is the standard deviation of the distribution.

### Neural dynamic range is linked to the temporal structure of brain activity

To this point, we have shown that the human connectome supports neural dynamics with a narrower distribution of dynamic ranges than the chimpanzee connectome (as well as other non-human primates). However, it remains unclear how dynamic range relates to the temporal structure of neural activity across the brain. Studies have shown that activity within brain regions exhibits a cortex-wide hierarchy of intrinsic neural timescales [25–27]. From these findings, we examine whether a brain region’s neural timescale may be related to its neural dynamic range. We extract the timescale by fitting the autocorrelation of the simulated neural activity with a single exponential decay (Fig. S14; see STAR Methods). We find that regional neural timescales (ranges: 0.12–0.24 s for humans and 0.12–0.55 s for chimpanzees) are significantly correlated with dynamic ranges, and this relation is stronger in chimpanzees (Fig. 4A; this finding also holds for macaques and marmosets as shown in Fig. S15A).

**Fig. 4.**
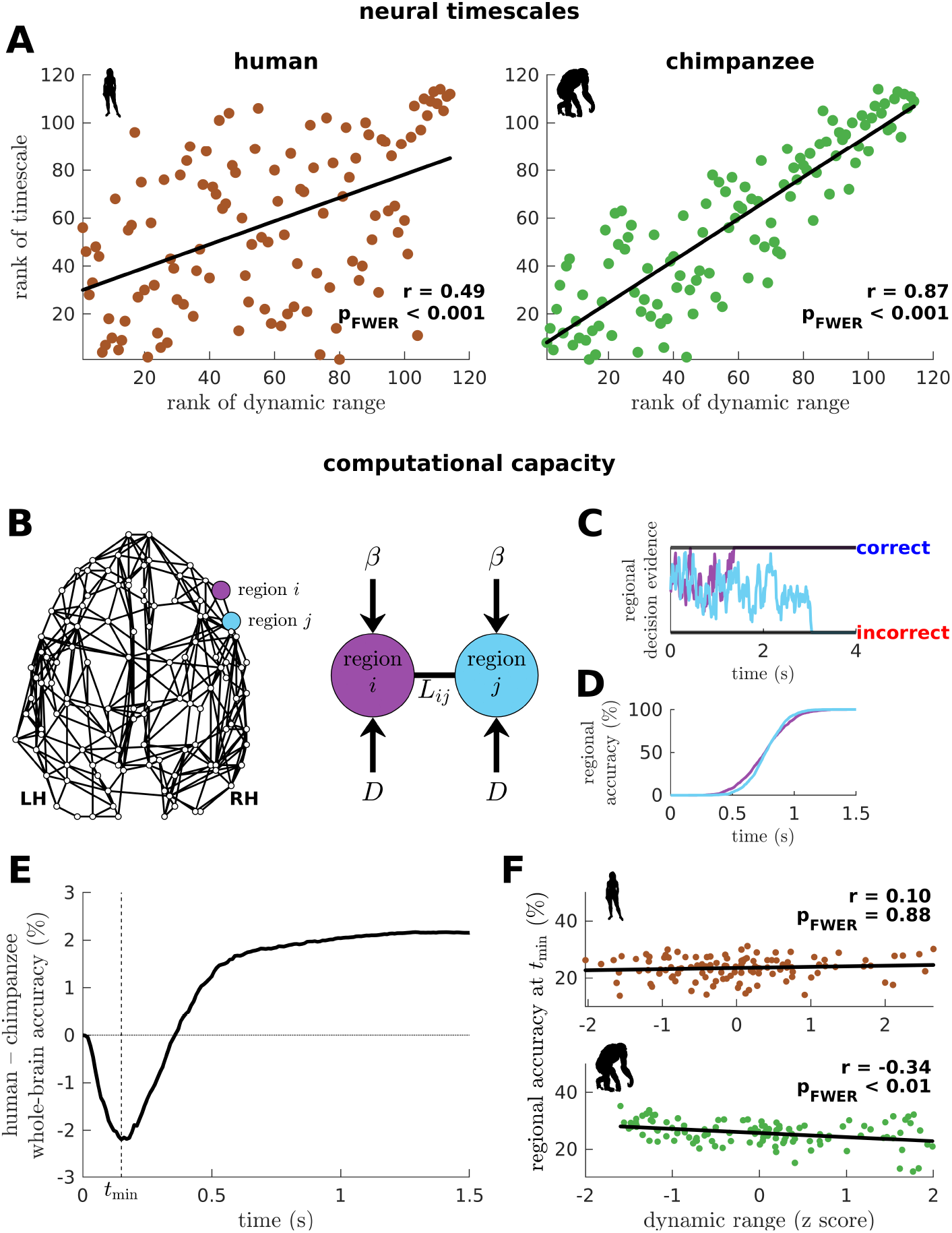
Human and chimpanzee neural timescales and their connectome’s computational capacity. (**A**) Ranked neural timescales as a function of ranked dynamic ranges. The solid line represents a linear fit with correlation coefficient r and FWER corrected p-value p_FWER_ after multiple comparisons. (**B**) Exemplar connectome and schematic diagram of the drift-diffusion model. In the model, each brain region accumulates decision evidence via a diffusion (Brownian) process with drift rate *β* and driving white noise with standard deviation *D*. Regions *i* and *j* are connected with Laplacian weight *L_ij_* based on the connectomic data. (**C**) Example time series of regional decision evidence across time for regions *i* and *j*, demonstrating how each region reaches a correct or incorrect decision. (**D**) Regional accuracy curves obtained by simulating the model for an ensemble of trials and calculating the rate of achieving the correct decision. (**E**) Difference in whole-brain accuracy across time between humans and chimpanzees. The dashed line shows the time (*t*_min_) at which the difference in accuracy between humans and chimpanzees is most negative (i.e., chimpanzee accuracy > human accuracy). (**F**) Regional accuracy at *t*_min_ (found in panel E) as a function of z-score-transformed dynamic ranges. The solid line represents a linear fit with correlation coefficient r and FWER corrected p-value p_FWER_ after multiple comparisons.

### Neural dynamic range affects the computational capacity of human and chimpanzee connectomes

We next ask what would be the implication of the differences in dynamic range distributions between humans and chimpanzees in terms of brain function. We hypothesize that these differences will likely impact the facilitation of whole-brain integration of neural processes, which has been found to be important for performing sensory-perceptual [28] and complex cognitive tasks [15] in humans. Note, however, that current whole-brain neuroimaging techniques cannot yet capture the direct effects of neural dynamic range on task performance. Moreover, new chimpanzee brain data via imaging or invasive recordings are not possible to be acquired for ethical reasons.

To provide insights into our question, we adopt a computational drift-diffusion model [29] (Fig. 4B), which is widely used to predict behavioral responses of both humans and animals performing tasks such as decision making. This model allows us to quantify the computational capacity of a brain region to achieve a decision threshold by integrating the evidence accumulated by its nearest neighbors. Here, we use the human and chimpanzee connectomes to define a region’s neighborhood. The model calculates the accumulation of evidence through time in each brain region via a noise-driven diffusion process until a set threshold is reached (Fig. 4C; there are two possible thresholds corresponding to a correct or incorrect decision). Then, we estimate each brain region’s accuracy in reaching a correct choice across an ensemble of trials (Fig. 4D) and average these values, representing the likely decision accuracy of the whole brain. At the end of our simulation, the human brain has a higher accuracy in achieving the correct choice compared to the chimpanzee brain (Fig. 4E; this finding also holds for macaques and marmosets as shown in Fig. S15B). Interestingly, we find that at earlier times (*t* < 0.36 s), the decision accuracy of the human brain is lower than the chimpanzee counterpart. This finding is driven by regions in the chimpanzee brain with low dynamic ranges that can reach correct decisions quickly (Fig. 4F).

### Testing of model predictions on empirical data

We have shown that a brain region’s neural dynamic range is tightly linked to the temporal structure of its activity (i.e., neural timescale), suggesting a role of dynamic range in local processing speeds. To test this prediction, we compare empirical cortical myelin maps of humans and chimpanzees [30] (Fig. 5A) to each region’s dynamic range. Myelin maps have been shown to be a good macroscale proxy of the cortical processing hierarchy in humans and non-human primates [30–32], where unimodal sensory-motor regions tend to be highly myelinated and transmodal regions lightly myelinated. Crucially, myelination has been found to be tightly coupled with electrophysiological measures of a region’s temporal processing speed [26]. Accordingly, we find that myelination is inversely related with neural dynamic range (Fig. 5B), with the relation stronger in chimpanzees. This result provides an empirical neurobiological support to our findings in Fig. 4A, demonstrating that dynamic range is related to a region’s processing speed.

**Fig. 5.**
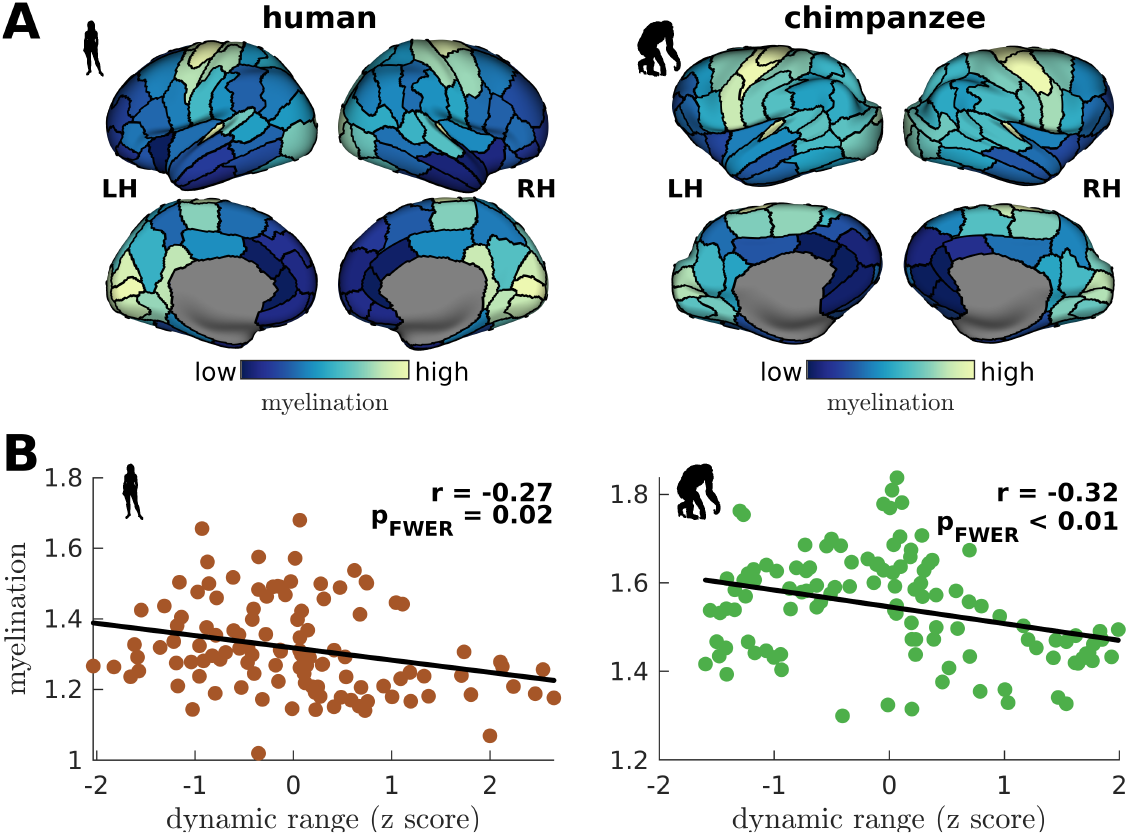
Testing of model predictions on myelination data. (**A**) Myelin maps visualized on inflated cortical surfaces. Light color represents high myelination and dark color represents low myelination. (**B**) Regional myelination as a function of z-score-transformed dynamic ranges. The solid line represents a linear fit with correlation coefficient r and FWER corrected p-value p_FWER_ after multiple comparisons.

We have also shown that the more constrained neural dynamics of the human brain compared to the chimpanzee brain allows better integration of whole-brain activity. We test this prediction on empirical neuroimaging (i.e., fMRI) data. Because there are no available chimpanzee fMRI data, we compare human with macaque data (the same data used in Fig. S1 to validate our model’s suitability). As predicted, we find that functional connectivity (FC) within large-scale networks is generally higher and more homogeneous in humans compared to macaques (Fig. 6A). The human brain also has higher between network functional connectivity, as demonstrated by a higher whole-brain *FC* (p_FWER_ < 0.001). Moreover, the human brain has lower functional path lengths than the macaque brain (Fig. 6B; similar metric used in Fig. S12 but applied here on functional connectivity matrices), which corresponds to better functional integration. We also estimated regional timescales by applying the same technique described in Fig. S14 on fMRI signals, finding that the human brain has faster timescales than the macaque brain (Fig. 6C). Overall, we have used available empirical human and non-human primate neuroimaging data to validate two key predictions of our model: neural dynamic range is linked to local processing speed and a narrower dynamic range distribution in humans allows better integration of whole-brain activity.

**Fig. 6.**
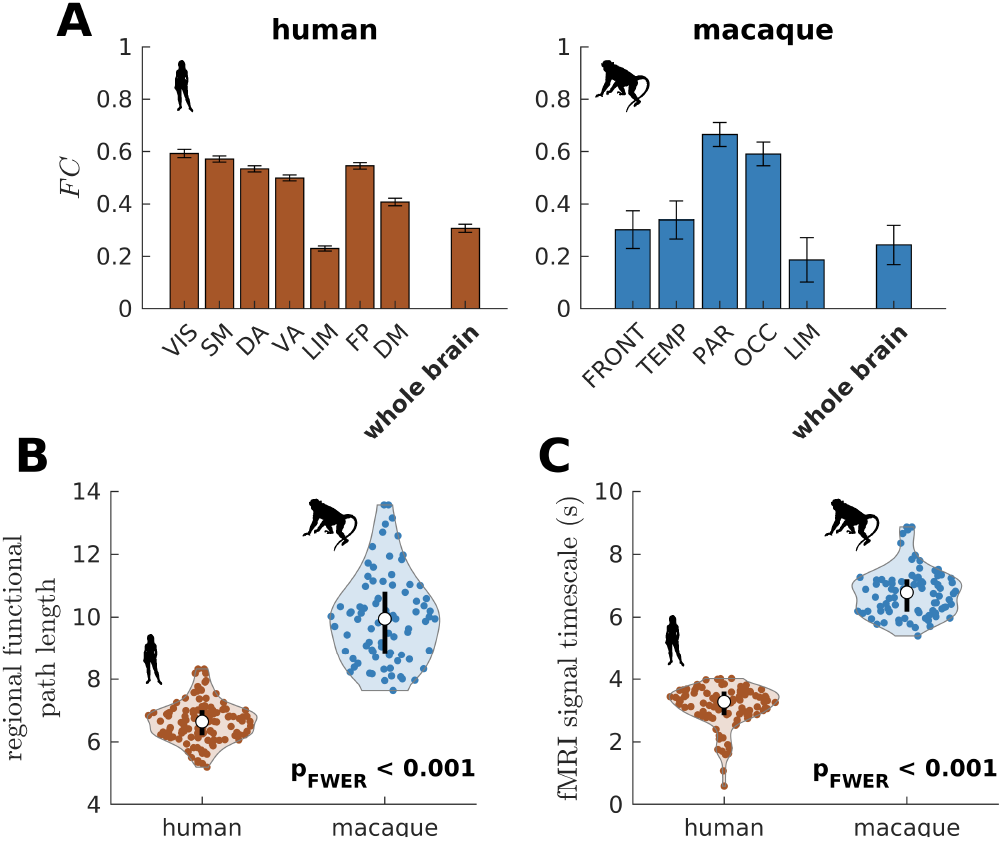
Testing of model predictions on functional neuroimaging data. (**A**) Functional connectivity (FC) within large-scale networks and across the whole brain of humans and macaques. The human large-scale networks are similar to those defined in Fig. S11. The macaque large-scale networks are: FRONT = Frontal; TEMP = Temporal; PAR = Parietal; OCC = Occipital; LIM = Limbic. The error bars are standard errors of the mean across subjects. (**B**) Violin plot of the distribution of regional functional path length across brain regions calculated from group-averaged functional connectivity matrices. Violin plot details are similar to those in Fig. 2B. (**C**) Violin plot of the distribution of fMRI signal timescales. Violin plot details are similar to those in Fig. 2B. For panels B and C, p is the FWER corrected p-value after multiple comparisons of the difference in the mean of the distribution between the species.

## DISCUSSION

There is converging evidence that the human brain, as compared to that of our primate relatives, has experienced significant structural reorganizations in association regions throughout evolution [3,33]. Here, we took advantage of a unique neuroimaging dataset of sex-matched and age-equivalent human and chimpanzee to study how the structural brain wirings of these species support different patterns of whole-brain neural dynamics underpinning brain function. Our results show that these differences determine how the activities of segregated regions are integrated across the brain, giving rise to distinct computational capacities of humans and chimpanzees.

For each species, we determined their brain regions’ response functions, which describe the regulation of intrinsic neural activity following brain-wide modulations in excitability. Modulations in global excitability could come from various sources, from external inputs [34] to internal influences such as neuromodulation [16–18]. The response function of each region encapsulates all these phenomena, differentiating it from typical input-output response curves [19,35], with its shape characterized by the neural dynamic range. A high dynamic range means that a region can slowly change its activity in response to a wide range of modulations in excitability. Conversely, a low dynamic range means that a region can quickly transition between low and high levels of activity within a narrow range of changes in excitability. We reasoned that the up or down regulation of local neural activity is crucial to facilitate communication across brain regions, enabling large-scale functional integration and related computations.

In humans, brain regions showed response functions that were more similar to one another than their chimpanzee counterparts. Moreover, the distribution of neural dynamic ranges in humans was narrower than that in chimpanzees and in other non-human primate species (i.e., macaques and marmosets). Thus, neural dynamic range seems to be a key unifying feature that sets human brains apart from other primate species. Note, however, that the generalization of the relationship between dynamic range and the evolutionary trajectory of non-human primates is beyond the scope of the current study and remains to be established. These results also propose the hypothesis that relatively subtle evolutionary differences in the connectomes of humans and chimpanzees [8] have a marked impact on large-scale neural dynamics supporting the ability of the brain to process information. This could be why human-specific features of connectome organization are vulnerable to brain dysfunction [10,36].

It has recently been shown that the structural, functional, and genetic properties of cortical regions are spatiotemporally organized in a hierarchical manner [27,32,37–41]. Accordingly, we found that neural dynamic ranges follow a dominant spatial gradient along the anterior-posterior brain axis, with anterior associative regions having more diffuse dynamic ranges than posterior sensory regions. This spatial gradient mirrors the cytoarchitectonic (e.g., cell density) organization observed in the brains of several mammalian species, including rodents and primates [42–44]. By analyzing the intrinsic timescales of fluctuations of neural activity [25–27], we showed that regional neural dynamic ranges correlated with regional neural timescales. Thus, dynamic range seems to capture local information processing speed, providing support to our prior hypothesis that brain regions with similar dynamic ranges are more likely to coactivate due to similar processing capacity. At the level of large-scale networks, chimpanzees showed a marked functional hierarchy in neural dynamic ranges compared to humans, with unimodal sensory-perceptual networks having lower values than transmodal associative networks. Importantly, this more pronounced hierarchy in dynamic ranges limits brain network integration in chimpanzees compared to humans. Human brain connectivity appears therefore to have evolved to support neural dynamics that maintain relatively high levels of integration between functionally segregated brain systems.

By using a computational drift-diffusion model [29], we assessed the functional consequences of neural dynamic range to the human and chimpanzee connectomes’ computational capacity. Over relatively long processing periods, the human connectome had a higher accuracy in achieving a correct choice compared to the chimpanzee connectome. However, over short periods of processing time, the human connectome performed worse than the chimpanzee counterpart. This latter result was attributed to the more heterogeneous distribution of dynamic ranges in chimpanzees, highlighting that diversity in local neural properties is important for rapid computations [35]. This regional diversity may explain why chimpanzees are able to perform at least as well as, or better than, humans in simple sensory-motor tasks [45]. Moreover, our findings provide a possible neural mechanism for why humans generally outperform chimpanzees in tasks requiring longer computational processing [46,47]. In line with our results, studies have suggested that behavioral differences between humans and chimpanzees are more prominent in complex tasks involving intersubjectivity (e.g., theory of mind [48]). This important brain capacity is known to be supported by the activity of the Default-Mode Network (DMN) [49,50], which displays significant genetic, anatomical, and functional differences between humans and non-human primates [22,51,52]. By making use of our simulations assessing computational capacity, we found that the accuracy of DMN for relatively long computations was more accurate in humans compared to chimpanzees (Fig. S16). This finding further suggests that the DMN may be critical in differentiating the functions of human and chimpanzee brains.

Results from the current study suggest that evolution has shaped the human brain to optimize fast transmodal integration of neural activity across brain regions supporting complex functions including social-cultural skills [40,53]. While our results are consistent with the hypothesis that the human brain has evolved to facilitate rapid associative computations [48], they also highlight that this evolutionary adaptation may hinder rapid processing within functionally specialized systems. The unique properties of human and chimpanzee brain dynamics may therefore be understood as an evolutionary tradeoff between functional segregation and integration. Collectively, our findings inform on the likely neural principles governing evolutionary shifts that could explain the differences in brain function between humans and our closest primate relatives [54].

## MATERIALS AND METHODS

### Connectomic data

#### Human and chimpanzee

Diffusion MRI data for 58 humans (*Homo sapiens*, 42.5 ± 9.8 years, female) and 22 chimpanzees (*Pan troglodytes*, 29.4 ± 12.8 years, female) were taken from previous studies [7,10]. Procedures were carried out in accordance with protocols approved by the Yerkes National Primate Research Center and the Emory University Institutional Animal Care and Use Committee (YER-2001206). All humans were recruited as healthy volunteers with no known neurological conditions and provided informed consent (IRB00000028). We only provide below a brief account of details of the data and we refer the readers to previous studies for further details. The diffusion MRI acquisition parameters for humans were: spin-echo echo planar imaging (EPI), isotropic voxel size of 2 mm, b-weighting of 1000 s/mm^2^, 8 bo-scans, and scan time of 20 mins. For chimpanzees: spin-echo EPI, isotropic voxel size of 1.8 mm, b-weighting of 1000 s/mm^2^, 40 bo-scans, and scan time of 60 mins. The acquired data were then preprocessed to correct for eddy-current, motion, susceptibility, and head motion distortions. Each participant’s cortex was then parcellated using a 114-area subdivision of the Desikan-Killiany atlas [55] (Table S1). Individual undirected connectome matrices were constructed via deterministic tractography to establish cortico-cortical connections between the 114 regions. In line with previous research [10], we removed idiosyncratic variations by taking the average weight across individuals of each connection that was consistently found in ≥60% of the individuals, resulting in a group-averaged weighted connectome for each species.

#### Macaque

The whole-brain macaque (*Macaca mulatta*) connectome was derived from 8 adult males using diffusion MRI. The data were taken from an open-source dataset, which provides in-depth description of the data [23]. In brief, the diffusion MRI acquisition parameters were: 2D EPI, isotropic voxel size of 1 mm, b-weighting of 1000 s/mm^2^, 64 directions, and 24 slices. The acquired data were then preprocessed to correct image distortion and to model fiber directions. Each macaque’s cortex was then parcellated into 82 regions following the Regional Map (RM) atlas [56]. Individual directed connectome matrices were constructed via probabilistic tractography to establish connections between the 82 regions and thresholded to remove weak connections (thresholds of 0% to 35%; see [23] for specific optimal threshold values used per individual). A group-averaged weighted connectome was obtained by taking the average weight of non-zero elements of the connectome matrices. It was then thresholded to match the connection density of the group-averaged chimpanzee connectome.

#### Marmoset

The marmoset (*Callithrix jacchus*) connectome data were downloaded from the Marmoset Brain Connectivity atlas (marmosetbrain.org) [24], which is a publicly available repository of cellular resolution cortico-cortical connectivity derived from neuroanatomical tracers. In brief, the connectome was reconstructed from 143 injections of six types of retrograde fluorescent tracers performed on 52 young adult marmosets (1.4–4.6 years, 21 females). Connection weights represent the fraction of labeled neurons found in the target area with respect to the total number of labeled neurons excluding the neurons in the injected area. The connections were projected onto the Paxinos stereotaxic atlas [57], comprising 116 cortical areas. Individual directed connectome matrices were constructed by including only areas with pairwise-complete connection values. Thus, the final connectome matrices were 55 × 55 in size. A group-averaged weighted connectome was obtained by taking the average weight of non-zero elements of the connectome matrices. It was then thresholded to match the connection density of the group-averaged chimpanzee connectome.

#### Human HCP

For replication of human results (Fig. S8), minimally preprocessed diffusion MRI data from 100 unrelated healthy young adult participants (29.1 ± 3.7 years, 54 females) were obtained from the Human Connectome Project (HCP) [20]. In brief, the diffusion MRI acquisition parameters were: isotropic voxel size of 1.25 mm, TR of 5520 ms, TE of 89.5 ms, b-weightings of 1000, 2000, and 3000 s/mm^2^, and 174 slices. The data were then preprocessed for bias-field correction and multi-shell multi-tissue constrained spherical deconvolution to model white matter, gray matter, and cerebrospinal fluid using the MRtrix software [58]. For each participant, tractograms were generated using 100 million probabilistic streamlines, anatomically-constrained tractography (ACT) [59], the 2^nd^-order Integration over Fiber Orientation Distributions algorithm (iFOD2), dynamic seeding [60], backtracking, streamline lengths of 5–250 mm, and spherical-deconvolution informed filtering of tractograms (SIFT2). Each participant’s tractogram was projected onto the cortex that was parcellated into 100 regions following the Schaefer atlas [61] to obtain the connectome matrices. A group-averaged weighted connectome was obtained by taking the average weight of non-zero elements of the connectome matrices. It was then thresholded to match the connection density of the chimpanzee connectome.

#### Macaque (CoCoMac)

For replication of macaque results (Fig. S13), a directed binary connectome was taken from an open-source dataset (the CoCoMac database) [62,63]. The connectome represents cortico-cortical structural connections between 71 regions derived from histological tract tracing studies.

### fMRI data

#### Human

The empirical human functional connectivity in Fig. S1B was derived from preprocessed functional MRI (fMRI) data of 100 unrelated healthy young adults from HCP (same participants used to calculate the HCP group-averaged connectome above) [20]. For each participant, functional connectivity (FC) was calculated by taking pairwise Pearson correlations of the BOLD-fMRI signal across 114 regions (Table S1). A group-averaged functional connectivity was obtained by taking the average of the individual functional connectivity matrices.

#### Macaque

The empirical macaque functional connectivity in Fig. S1B was derived from preprocessed fMRI data of 8 adult rhesus macaques (same subjects used to calculate the macaque group-averaged connectome above) [23]. For each subject, *FC* was calculated by taking pairwise Pearson correlations of the BOLD-fMRI signal across 82 regions [56]. A group-averaged functional connectivity was obtained by taking the average of the individual functional connectivity matrices.

### Cortical myelination data

Human and chimpanzee cortical myelination data were obtained from [30]. The data represent average myelination maps estimated from T1w:T2w ratio values. The maps were then parcellated using our 114-region atlas (Table S1).

### Cortical surfaces

For visualization purposes, we mapped some results into template human and chimpanzee inflated cortical surfaces (Figs. 2C, 5A, S11, S12C, and S16) obtained from https://balsa.wustl.edu/study/Klr0B, which is a public repository of data from [30].

### Computational models

#### Reduced Wong-Wang neural model

To simulate local neural dynamics on the connectome, we used the reduced Wong-Wang biophysical model, also known as the dynamic mean field model, which is an established model derived from a mean-field spiking neuronal network [11–14]. Each brain region *i* is governed by the following nonlinear stochastic differential equation:

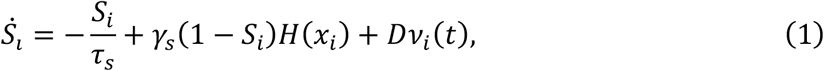

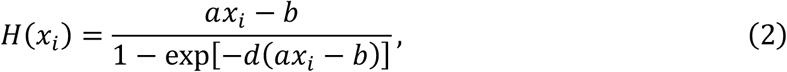

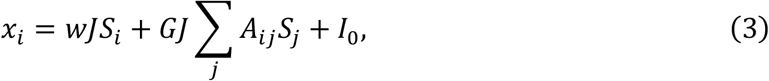

where *S_i_*, *H*(*x_i_*), and *x_i_* represent the synaptic gating variable, firing rate, and total input current, respectively. The synaptic gating variable *S_i_* in Eq. (1) is governed by the time constant *τ_s_* = 0.1 s, saturation rate *γ_s_* = 0.641, firing rate *H*, and independent zero-mean Gaussian noise *v_i_* with standard deviation *D* = 0.003. The firing rate *H* is a nonlinear input-output function defined in Eq. (2) governed by the total input current *x_i_* with constants *a* = 270 (V nC)^-1^, *b* = 109 Hz, and *d* = 0.154 s. The total input current *x_i_* is determined in Eq. (3) by the recurrent connection strength *w*, synaptic coupling *J* = 0.2609 nA, global scaling constant *G* = 0.2, connection strength *A_ij_* between regions *i* and *j*, and excitatory subcortical input *I*_0_ = 0.33 nA. The parameter values were taken from previous works [13,14]. Note that the value of the global scaling constant was fixed for all species. This is to ensure that we can directly compare variations in neural dynamics. We simulated the model by numerically solving Eq. (1) using the Euler-Maruyama scheme for a time period of 720 s and a time step of 0.01 s. We then calculated the time-average value of the synaptic gating variable *S* after removing transients, which we used to represent neural dynamics in our analyses.

#### Balloon-Windkessel hemodynamic model

To obtain the simulated functional connectivity in Figs. 2D and S1, we fed the neural activity *S_i_* from Eq. (1) to the Balloon-Windkessel hemodynamic model, which is an established model for simulating BOLD-fMRI signals [64]. Each brain region *i* is governed by the following nonlinear differential equations:

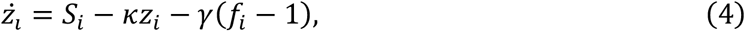

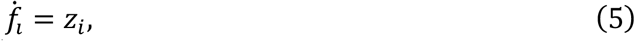

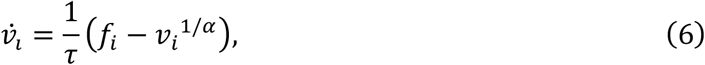

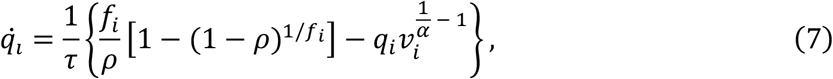

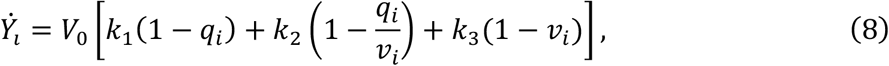

where *z_i_*, *f_i_*, *v_i_*, *q_i_*, and *Y_i_* represent the vasodilatory signal, blood inflow, blood volume, deoxyhemoglobin content, and BOLD-fMRI signal, respectively. The model parameters and their values are defined as follows: signal decay rate *κ* = 0.65 s^-1^, elimination rate *γ* = 0.41 s^-1^, hemodynamic transit time *τ* = 0.98 s, Grubb’s exponent *α* = 0.32, resting oxygen extraction fraction *ρ* = 0.34, resting blood volume fraction *V*_0_ = 0.02, and fMRI parameters *k*_1_ = 4.10, *k*_2_ = 0.58, *k*_3_ = 0.53. The parameter values were taken from previous works [64]. We simulated the model for a time period of 720 s and the time series were downsampled to a temporal resolution of 0.72 s to match the resolution of typical empirical BOLD-fMRI signals. Functional connectivity (FC) was calculated by taking pairwise Pearson correlations of *Y_i_* (after removing transients) across all regions. The within-network functional connectivity in Fig. 2D was obtained by taking the average of the *FC* between regions comprising each network.

#### Wilson-Cowan neural model

To show that our results generalize beyond our choice of biophysical model, we also simulated local neural dynamics using the Wilson-Cowan model [65] (Fig. S9). We chose this model because of its known ability to reproduce diverse large-scale neural phenomena [66]. Each brain region *i* comprises interacting populations of excitatory (*E*) and inhibitory (*I*) neurons governed by the following nonlinear stochastic differential equations:

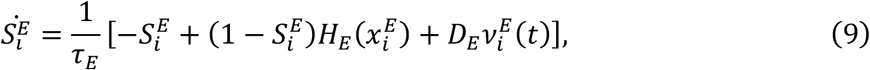

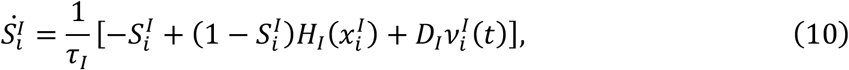

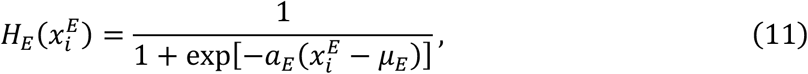

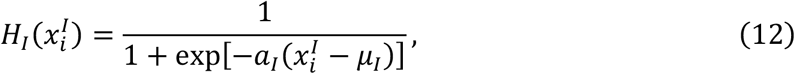

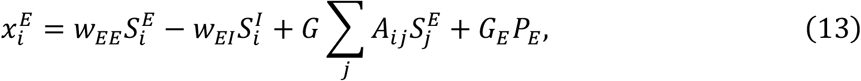

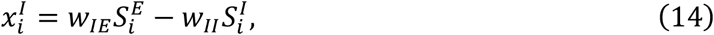

where *S_i_*, *H*(*x_i_*), and *x_i_* represent the firing rate, non-linear activation function, and weighted sum of firing rates, respectively, for *E* and *I* populations. The dynamics of the firing rates 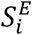 and 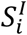 in Eqs. (9) and (10) are parameterized by the excitatory time constant *τ_E_* = 2.5×10^-3^ s, inhibitory time constant *τ_I_* = 3.75×10^-3^ s, activation functions *H_E_* and *H_I_*, and independent zero-mean Gaussian noise 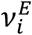 and 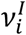 with standard deviations *D_E_* = 5×10^-5^ and *D_E_* = 5×10^-5^, respectively. The activation functions *H_E_* and *H_I_* in Eqs. (11) and (12) are defined by sigmoids parameterized by the gain constants *α_E_* = 1.5 and *α_I_* = 1.5 and firing thresholds *μ_E_* = 3 and 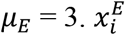 and 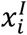 are determined in Eqs. (13) and (14) by the excitatory-excitatory recurrent connection strength *w_EE_* = 16, excitatory-inhibitory connection strength *w_EI_* = 12, inhibitory-excitatory connection strength *w_IE_* = 15, inhibitory-inhibitory recurrent connection strength *w_II_* = 3, global scaling constant *G* = 2, connection strength *A_ij_* between regions *i* and *j*, and excitatory drive *P_E_* = 1 scaled by *G_E_* = 0.5. The parameter values were taken from previous works [65,66]. We simulated the model by numerically solving Eqs. (9) and (10) using the Euler-Maruyama scheme for a time period of 15 s and a time step of 0.001 s. We then calculated the time-average value of the excitatory firing rate *S^E^* after removing transients, which we used to represent neural dynamics in our analyses.

#### Drift-diffusion model

To simulate the ability of human and chimpanzee connectomes to reach a binary decision, we implemented a computational drift-diffusion model [29] (Fig. 4B). Each brain region *i* is governed by the following stochastic differential equation:

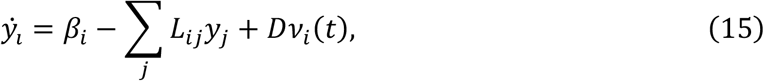

where *y_i_* is the evidence at time *t*, *β_i_* is the drift rate, *L_ij_* is the Laplacian weight of the connection between regions *i* and *j*, and *v_i_* is an independent zero-mean Gaussian noise with standard deviation *D*. The Laplacian matrix *L* is obtained via 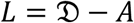, where *A* is the connectivity matrix and 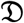 is a diagonal matrix of node strengths such that the *i*th diagonal element is ∑*_j_ A_ij_*. To focus on the contribution of the connectome itself, we fixed the drift rates of the regions to *β_i_* = 1 and *D* = 1. We verified that changing these parameter values did not change the results of the study. Through the simple diffusion process implemented by the model, each region accumulates the decision evidence through time until it reaches a boundary threshold *θ* = ±1 where a decision is said to be reached. Without loss of generality, we assumed *θ* = 1 to be the correct decision (Fig. 4C). We simulated the model by numerically solving Eq. (15) using the Euler-Maruyama scheme for a time period of 5 s and a time step of 0.01 s. We then calculated the decision accuracy versus time of each region across an ensemble of 1000 trials (Fig. 4D).

### Measures of neural dynamics properties

#### Neural dynamic range

For each brain region, we analyzed its response function reflecting how activity changes versus global modulations in the strength of recurrent connections (i.e., *S* versus *w* for the reduced Wong-Wang model and *S^E^* versus *w_EE_* for the Wilson-Cowan model). The response function was characterized in terms of the neural dynamic range, mathematically defined as:

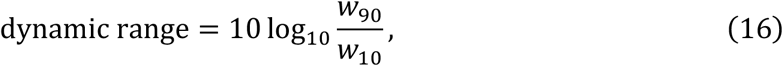

where *w_x_* is the corresponding global recurrent strength at *S_x_* with *x* = {10,90} and

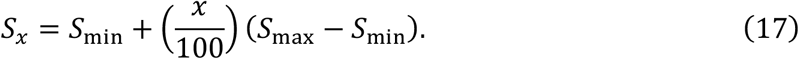

We then pooled together the dynamic ranges of brain regions for each species of interest to create a distribution with standard deviation σ.

#### Neural timescale

To estimate neural timescales for each brain region, we simulated neural activity via the model described in Eqs. (1) to (3). We used a global recurrent strength of *w* = 0.45 to produce neural dynamics in a biologically plausible regime; i.e., dynamics with relatively low firing rate and not fully synchronized [67]. Following recent studies [25,26], we quantified the neural timescale of each brain region *i* by fitting the autocorrelation of *S_i_*(*t*) with a single exponential decay function (via non-linear least-squares) with form

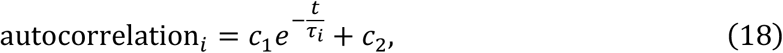

where *c*_1_ and *c*_2_ are fitting constants and *τ_i_* is the estimated neural timescale (Fig. S14). We verified that fitting with a double exponential decay function did not change the results of the study.

### Additional confirmatory analyses

We performed several confirmatory analyses to check the robustness of our results. In particular, we addressed potential effects of differences in individual-specific connectomes (Fig. S2), differences in connection density (Fig. S3), variability of connection strengths across participants (Fig. S4), differences in human and chimpanzee data sample size (Fig. S5), and existence of activity propagation delays between brain regions (Fig. S6). The procedures for each confirmatory analysis are described below.

#### Individual-specific analysis

Brain dynamics resulting from group-averaged connectomes could differ when individual-specific connectomes are used. Hence, we repeated the analysis in Fig. 2 to the connectome of each human and chimpanzee participant to produce Fig. S2.

#### Connection density

The resulting connection densities of the group-averaged human and chimpanzee connectomes were different (13.7% versus 11.6%, respectively). Hence, we created a new human connectome by pruning weak connections such that its density matches the density of the chimpanzee connectome. We then repeated the analysis in Fig. 2 to produce Fig. S3.

#### Inter-individual variability of connection strengths

The quality of connectomic data across participants in each species may be different due to potential additional confounds unable to be corrected by the implemented data preprocessing methods. Thus, within each species, we calculated the variability of weights of each connection *A_ij_* across participants (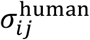 and 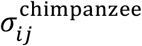). We then rescaled the human and chimpanzee connectomes to match each other’s inter-individual variability. Specifically, we multiplied the human connectome weights by 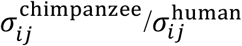 and the chimpanzee connectome weights by 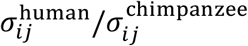. We then repeated the analysis in Fig. 2 to produce Fig. S4.

#### Sample sizes

The sample sizes of the human and chimpanzee connectomic data differ (N = 58 for humans and N = 22 for chimpanzees). Hence, we randomly sampled 22 human participants to match the sample size of the chimpanzee data and calculated the corresponding new group-averaged human connectome. The resampling procedure was repeated for 100 trials. For each trial, we repeated the analysis in Fig. 2 to produce Fig. S5.

#### Activity propagation delay

The model we used, as described in Eqs. (1) to (3), assumes that the activity of a brain region propagates and affects instantaneously the activity of all other regions connected to it (i.e., propagation time delay is zero). However, due to the spatial embedding of the brain and the finiteness of activity propagation speeds, we modified the second term of Eq. (3) to incorporate non-zero time delays as follows:

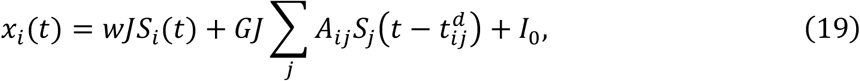

where 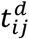 is the propagation time delay between regions *i* and *j*. We approximated the time delays as 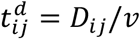, where *D_ij_* is the Euclidean distance between the centroids of regions *i* and *j* and *v* is the propagation speed. We assumed *v* to be a constant with value 10 m/s [68] and was the same for both human and chimpanzee brains; choosing other values of *v* did not change our results. Meanwhile, *D_ij_* is specific to human and chimpanzee brains to account for differences in brain sizes. In our analysis, we obtained *D_ij_* from one randomly chosen representative human and chimpanzee; the resulting distributions of time delays are shown in Fig. S6A. We then repeated the analysis in Fig. 2 to produce Figs. S6B and S6C.

#### Heterogeneous excitatory input

The model we used, as described in Eqs. (1) to (3), assumes that the activities of brain regions are driven by a constant excitatory input *I*_0_. We tested whether varying *I*_0_ per region affects the results of the study. In particular, we incorporated a gradient of excitatory input across the anatomical cortical hierarchy, with unimodal regions having higher input and transmodal regions having lower input [14]. Inspired by previous works using a region’s total connection strength as a proxy of the cortical hierarchy [69,70], we modified the last term of Eq. (3) as follows:

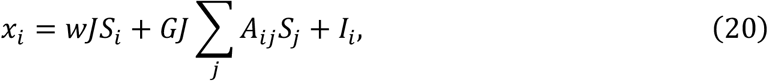

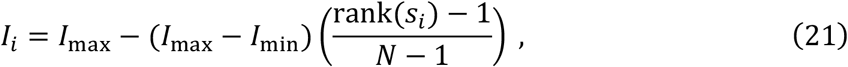

where *I*_max_ = 0.33 nA, *I*_min_ = 0.28 nA, *s_i_* = ∑*_j_ A_ij_* is the total connection strength of region *i*, and *N* is the total number of regions. The resulting excitatory input of regions in human and chimpanzee brains are shown in Fig. S7A. We then repeated the analysis in Fig. 2 to produce Figs. S7B and S7C. Note that the values of *I*_max_ and *I*_min_ were chosen to match the spread of estimated excitatory inputs found in [14]. However, we verified that using other values of *I*_max_ and *I*_min_ did not change the results of the study.

### Additional graph theoretical analysis

To bridge our functional results with the structural property of the connectome, we performed an additional analysis using concepts in graph theory [71]. In particular, we quantified the level of integration imposed by the connectome, which is well captured by the measure of path length (low path length = high integration; Fig. S12). Path length corresponds to the total topological distance of the shortest path between two regions. For our weighted connectomes, we defined the topological distance to be inversely proportional to the weight of connection (i.e., distance = 1/weight). Then, for each brain region, the regional path length was calculated by taking the average of the path lengths between a region and all other regions.

## DATA AND CODE AVAILABILITY

All source data and MATLAB codes to perform sample simulations, analyze results, and generate the main and supplementary figures of this study are openly available at https://github.com/jchrispang/evolution-brain-tuning.

## ACKNOWLEDGMENTS

We thank John Murray, Alex Fornito, and Luke Hearne for valuable scientific discussions. Human HCP data used for replication of results were provided by the Human Connectome Project, Wu-Minn Consortium (Principal Investigators: David Van Essen and Kamil Ugurbil; 1U54MH091657) funded by the 16 NIH Institutes and Centers that support the NIH Blueprint for Neuroscience Research, and by the McDonnell Center for Systems Neuroscience at Washington University. This work was supported by National Health and Medical Research Council grants 11144936 and 1145168 to J.A.R, Netherlands Organization for Scientific Research grants ALWOP.179 and VIDI (452-16-015) to M.P.V.D.H., and National Health and Medical Research Council grants 1138711 and 2001283 to L.C.

## AUTHOR CONTRIBUTIONS

Conceptualization: J.C.P., J.A.R., M.P.V.D.H., L.C.; Methodology: J.C.P., J.A.R., M.P.V.D.H., L.C.; Software: J.C.P.; Formal analysis: J.C.P.; Investigation: J.C.P., J.K.R., J.A.R., M.P.V.D.H., L.C.; Resources: J.A.R., M.P.V.D.H.; Visualization: J.C.P., J.A.R., M.P.V.D.H., L.C.; Funding acquisition: J.A.R., M.P.V.D.H., L.C.; Supervision: J.A.R., L.C.; Writing – original draft: J.C.P., L.C.; Writing – review & editing: J.C.P., J.K.R., J.A.R., M.P.V.D.H., L.C.

## COMPETING INTERESTS

The authors declare no competing interests.

## SUPPLEMENTARY INFORMATION

**Fig. S1.**
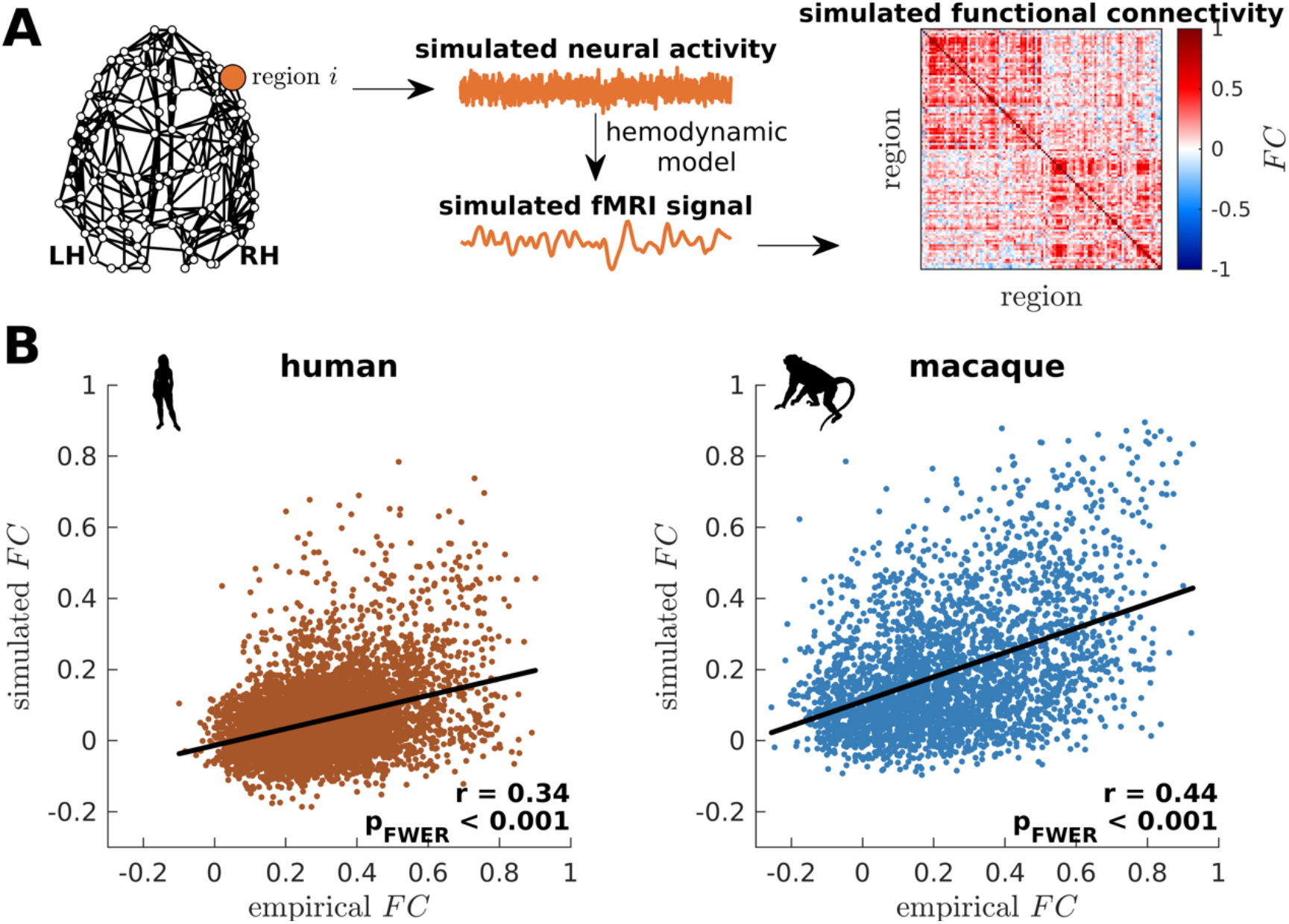
Validation of simulated dynamics on empirical functional neuroimaging data. (**A**) From the connectome, neural activity is simulated using the model presented in Fig. 1B. This activity is fed into a hemodynamic model to obtain a simulated fMRI signal for each brain region. Finally, functional connectivity (FC) is calculated by taking pairwise Pearson correlations of the simulated fMRI signals across all regions. (**B**) Association between simulated and empirical functional connectivity for humans and macaques. The simulated functional connectivity is calculated using model parameters that optimize model-data fitting. Each dot represents the pairwise *FC*. The solid line represents a linear fit with correlation coefficient r and FWER corrected p-value p_FWER_ after multiple comparisons.

**Fig. S2.**
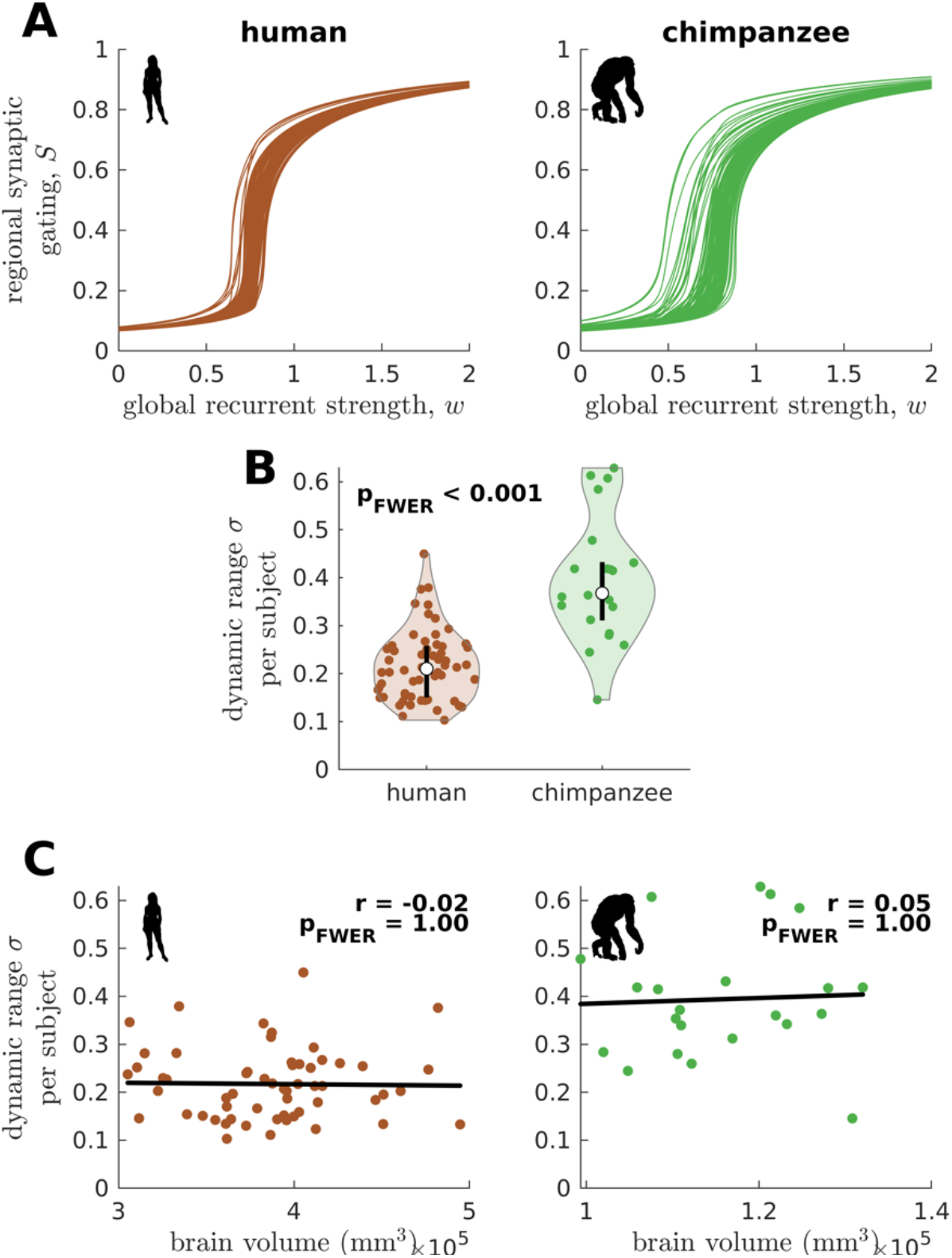
Confirmatory analysis on individual-specific connectomes and accounting for total brain volume. (**A**) Regional neural dynamics as a function of global recurrent strength (*w*) for exemplar human and chimpanzee participants. (**B**) Violin plot of the standard deviation (*σ*) of the distribution of dynamic ranges across brain regions for each participant. Each violin shows the first to third quartile range (black line), median (white circle), raw data (dots), and kernel density estimate (outline). p is the FWER corrected p-value after multiple comparisons of the difference in the mean of the distribution between the species. (**C**) Dynamic range standard deviation (*σ*) for each participant as a function of total brain volume. The solid line represents a linear fit with correlation coefficient r and FWER corrected p-value p_FWER_ after multiple comparisons.

**Fig. S3.**
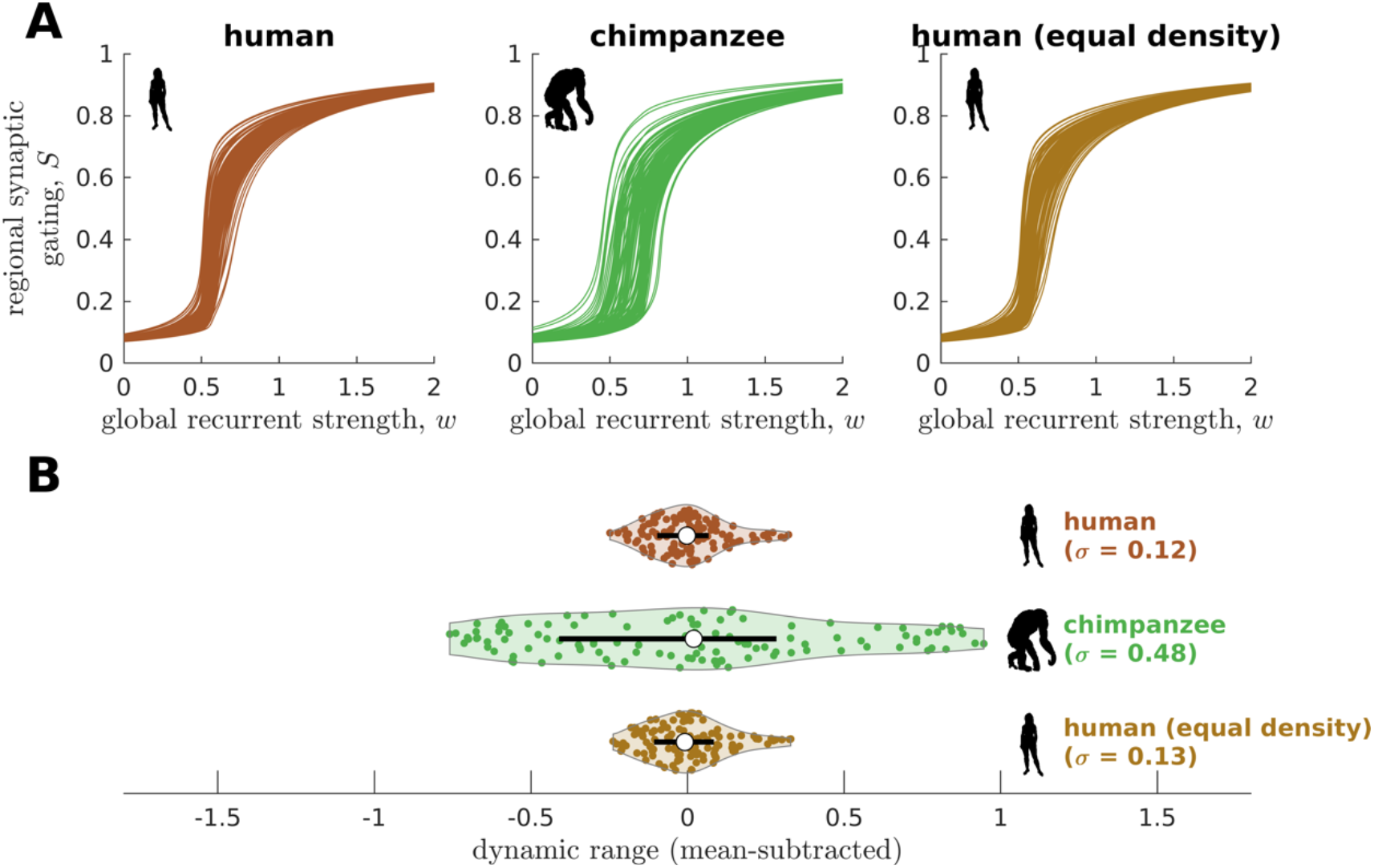
Confirmatory analysis on human and chimpanzee connectomes of equal connection density. (**A**) Regional neural dynamics as a function of global recurrent strength (w) for the original human connectome, original chimpanzee connectome, and human connectome pruned to have an equal density as the original chimpanzee connectome. (**B**) Violin plot of the distribution of dynamic ranges across brain regions. Each violin shows the first to third quartile range (black line), median (white circle), raw data (dots), and kernel density estimate (outline). The data are mean-subtracted for visual purposes. σ is the standard deviation of the distribution.

**Fig. S4.**
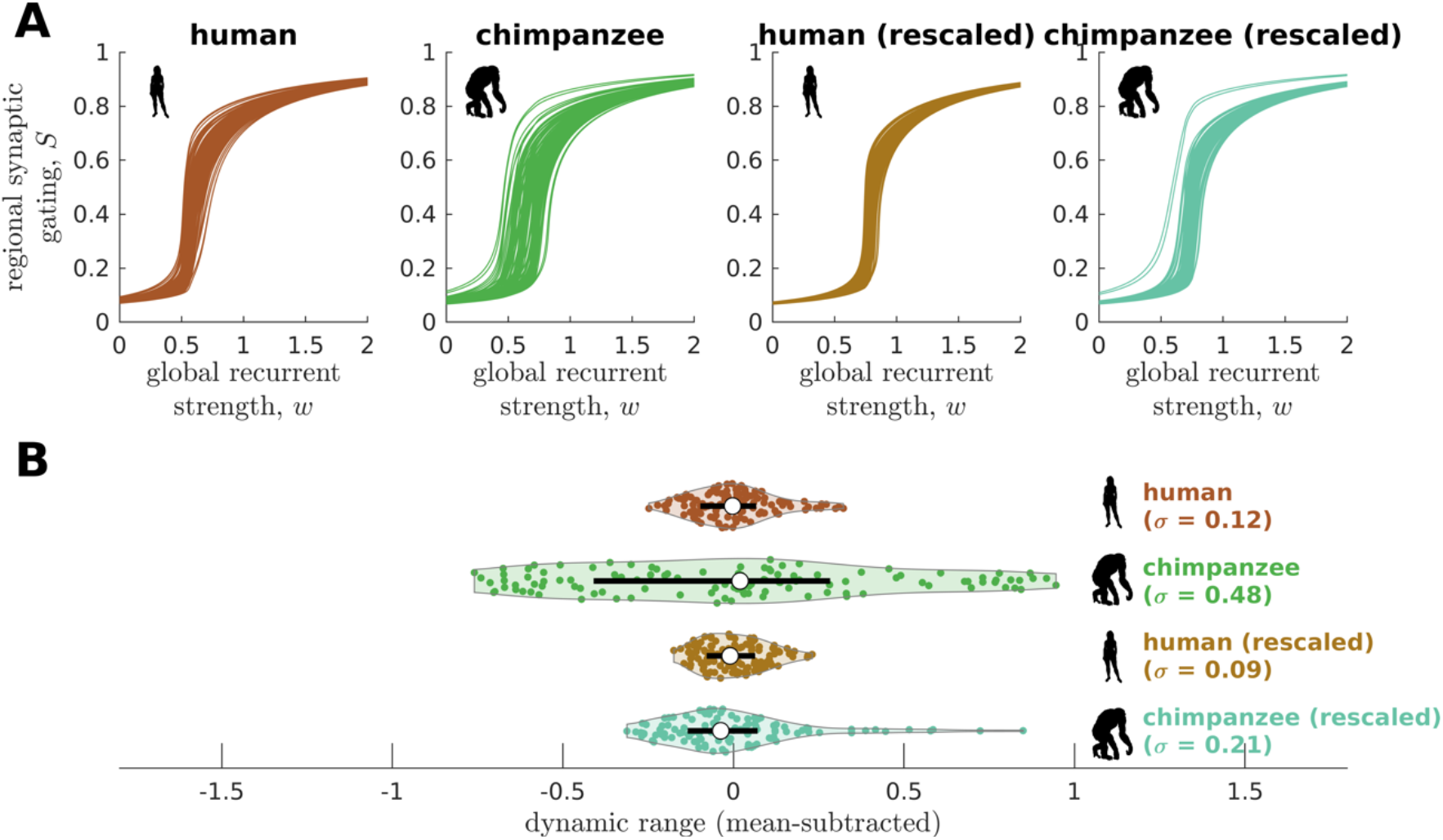
Confirmatory analysis accounting for inter-individual variability of connectomic data. (**A**) Regional neural dynamics as a function of global recurrent strength (*w*) for the original human connectome, original chimpanzee connectome, human connectome rescaled to match the inter-individual variability of the original chimpanzee connectome, and chimpanzee connectome rescaled to match the inter-individual variability of the original human connectome. (**B**) Violin plot of the distribution of dynamic ranges across brain regions. Each violin shows the first to third quartile range (black line), median (white circle), raw data (dots), and kernel density estimate (outline). The data are mean-subtracted for visual purposes. σ is the standard deviation of the distribution.

**Fig. S5.**
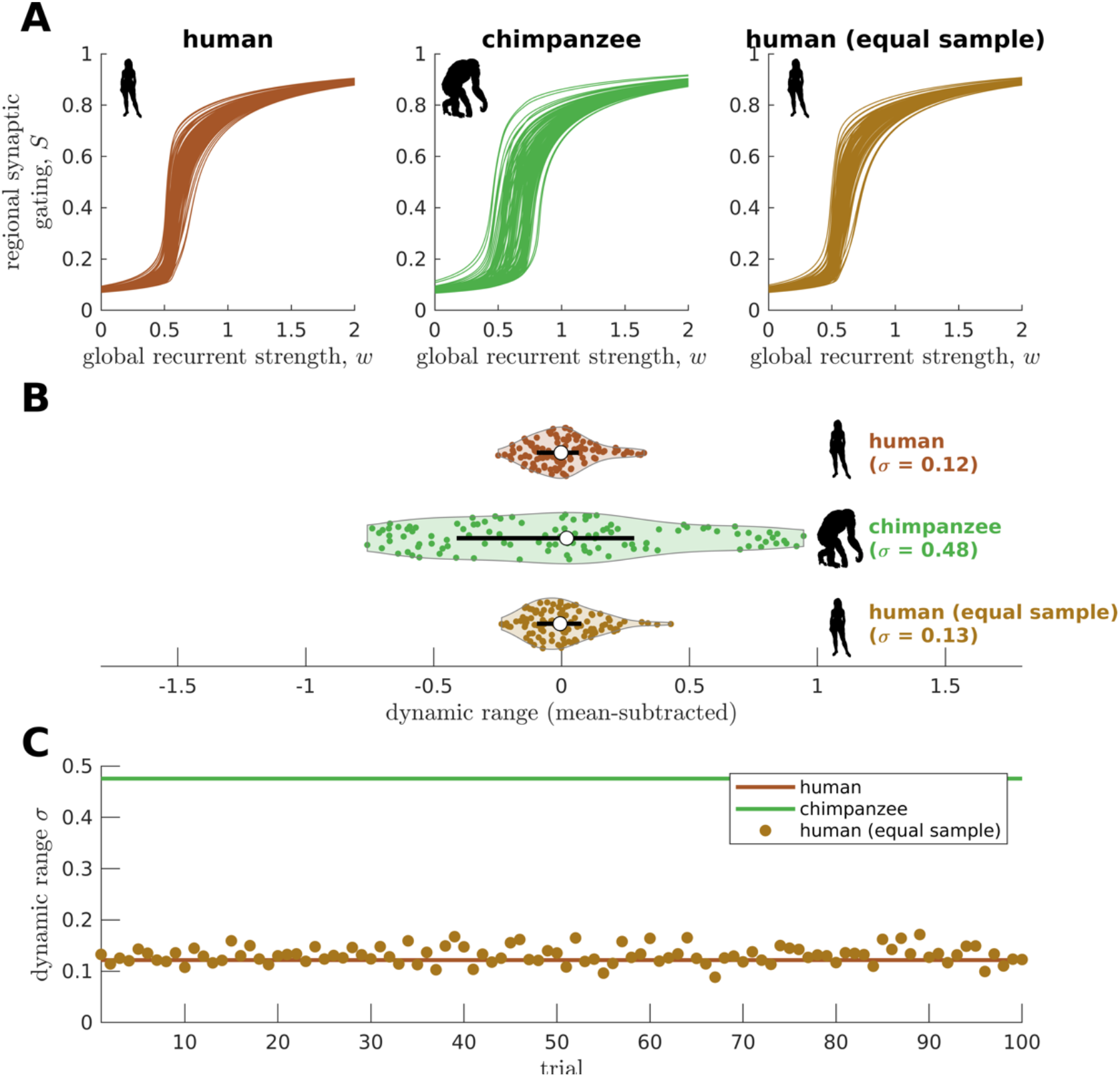
Confirmatory analysis on matched sample size. (**A**) Regional neural dynamics as a function of global recurrent strength *w*) for the original human connectome, original chimpanzee connectome, and an exemplar human connectome averaged from a sample of random human participants of the same size as the chimpanzee group (N = 22). (**B**) Violin plot of the distribution of dynamic ranges across brain regions. Each violin shows the first to third quartile range (black line), median (white circle), raw data (dots), and kernel density estimate (outline). The data are mean-subtracted for visual purposes. *σ* is the standard deviation of the distribution. (**C**) Dynamic range standard deviation (*σ*) for multiple random sampling trials of human participants. The solid lines represent the results for the original human and chimpanzee connectomes.

**Fig. S6.**
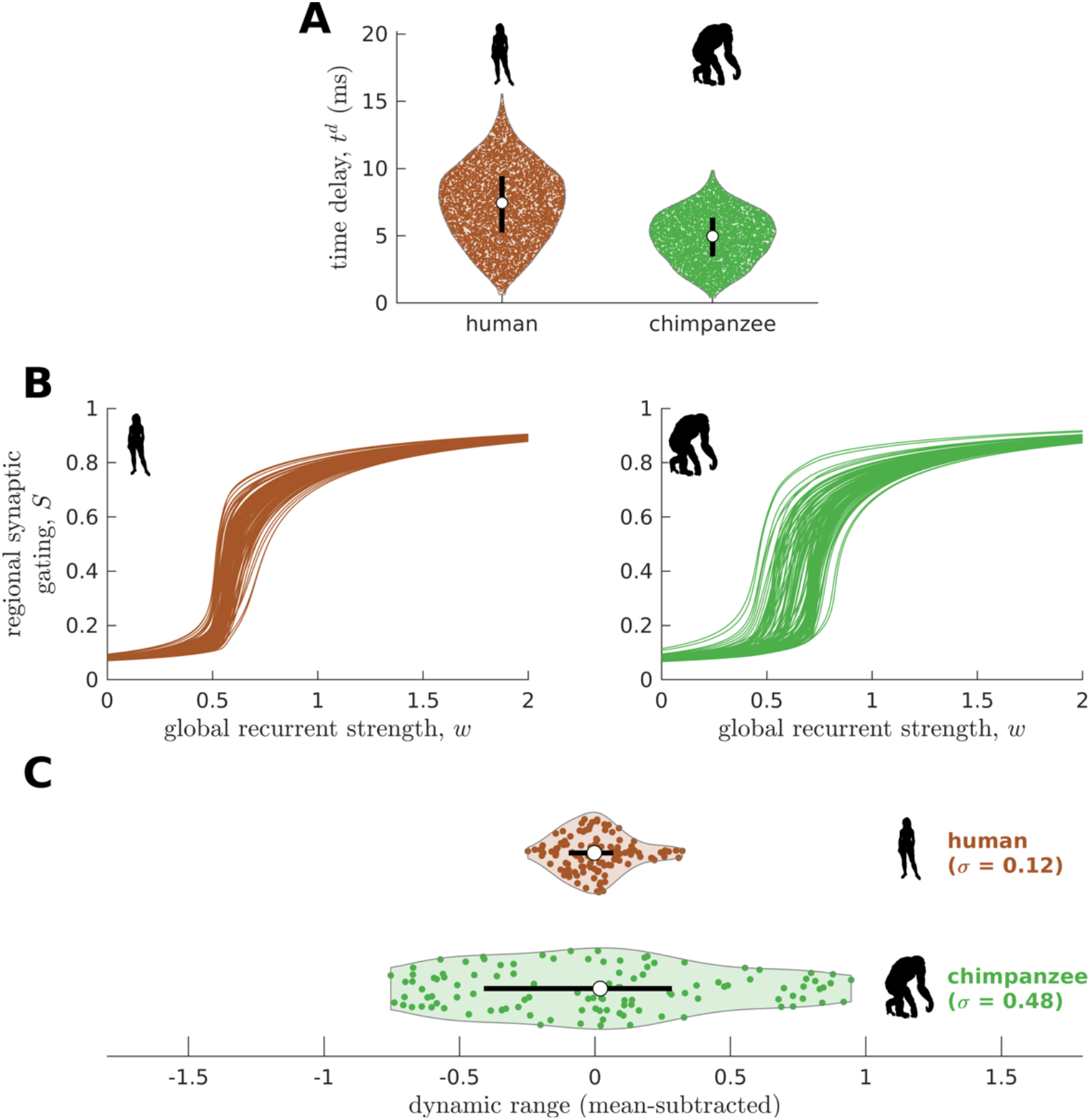
Confirmatory analysis accounting for activity propagation delays between brain regions. (**A**) Violin plot of the distribution of propagation time delays (*t^d^*) across all connections for a representative human and chimpanzee. Each violin shows the first to third quartile range (black line), median (white circle), raw data (dots), and kernel density estimate (outline). (**B**) Regional neural dynamics as a function of global recurrent strength (*w*). (**C**) Violin plot of the distribution of dynamic ranges across brain regions. Each violin shows the first to third quartile range (black line), median (white circle), raw data (dots), and kernel density estimate (outline). The data are mean-subtracted for visual purposes. *σ* is the standard deviation of the distribution.

**Fig. S7.**
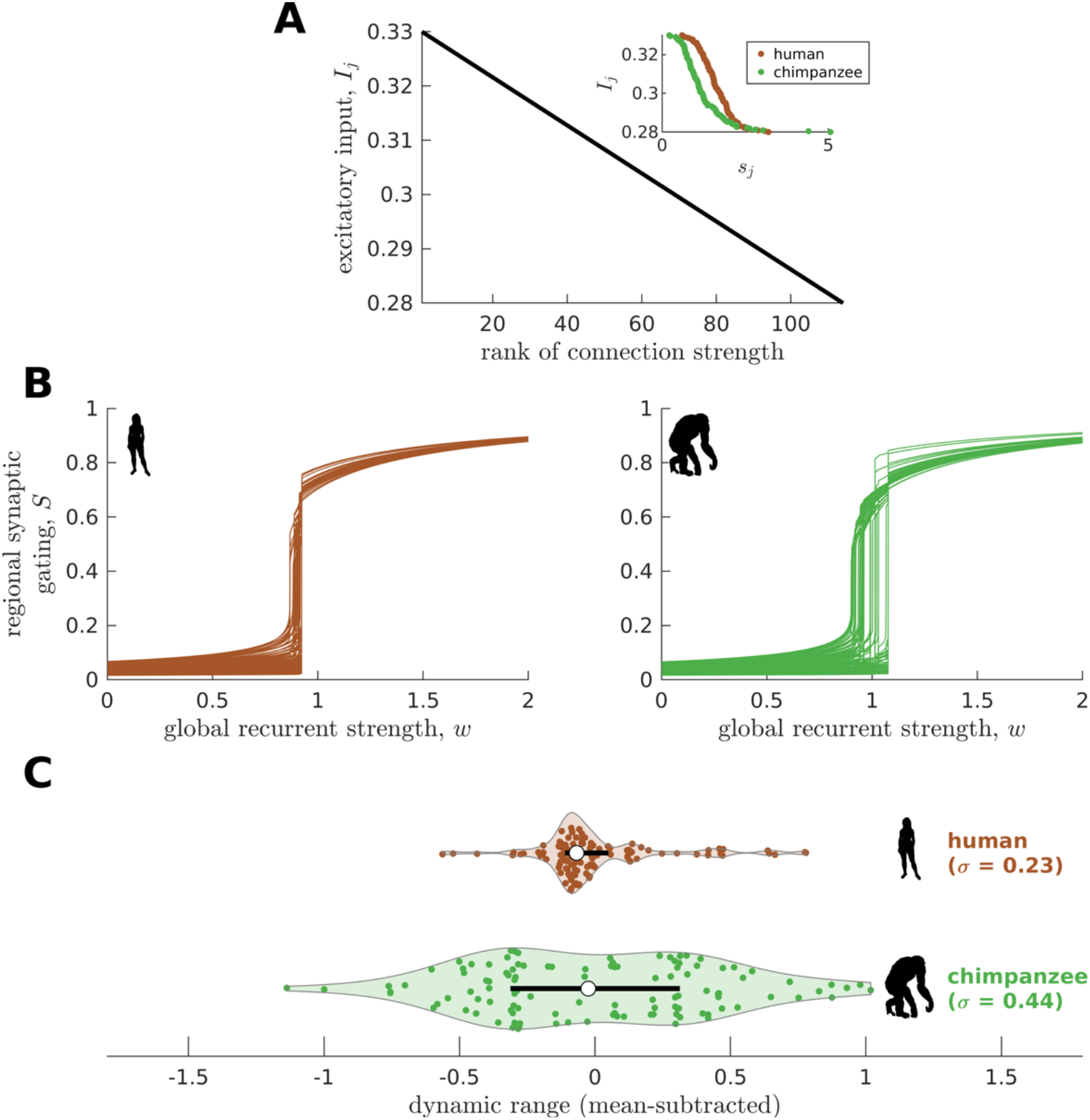
Confirmatory analysis accounting for heterogeneous excitatory input across brain regions. (**A**) The excitatory input in each brain region *I_j_* is inversely proportional to the rank of its total connection strength (*s_j_*). The inset shows the actual relationship between *I_j_* and *s_j_*. (**B**) Regional neural dynamics as a function of global recurrent strength (*w*). (**C**) Violin plot of the distribution of dynamic ranges across brain regions. Each violin shows the first to third quartile range (black line), median (white circle), raw data (dots), and kernel density estimate (outline). The data are mean-subtracted for visual purposes. *σ* is the standard deviation of the distribution.

**Fig. S8.**
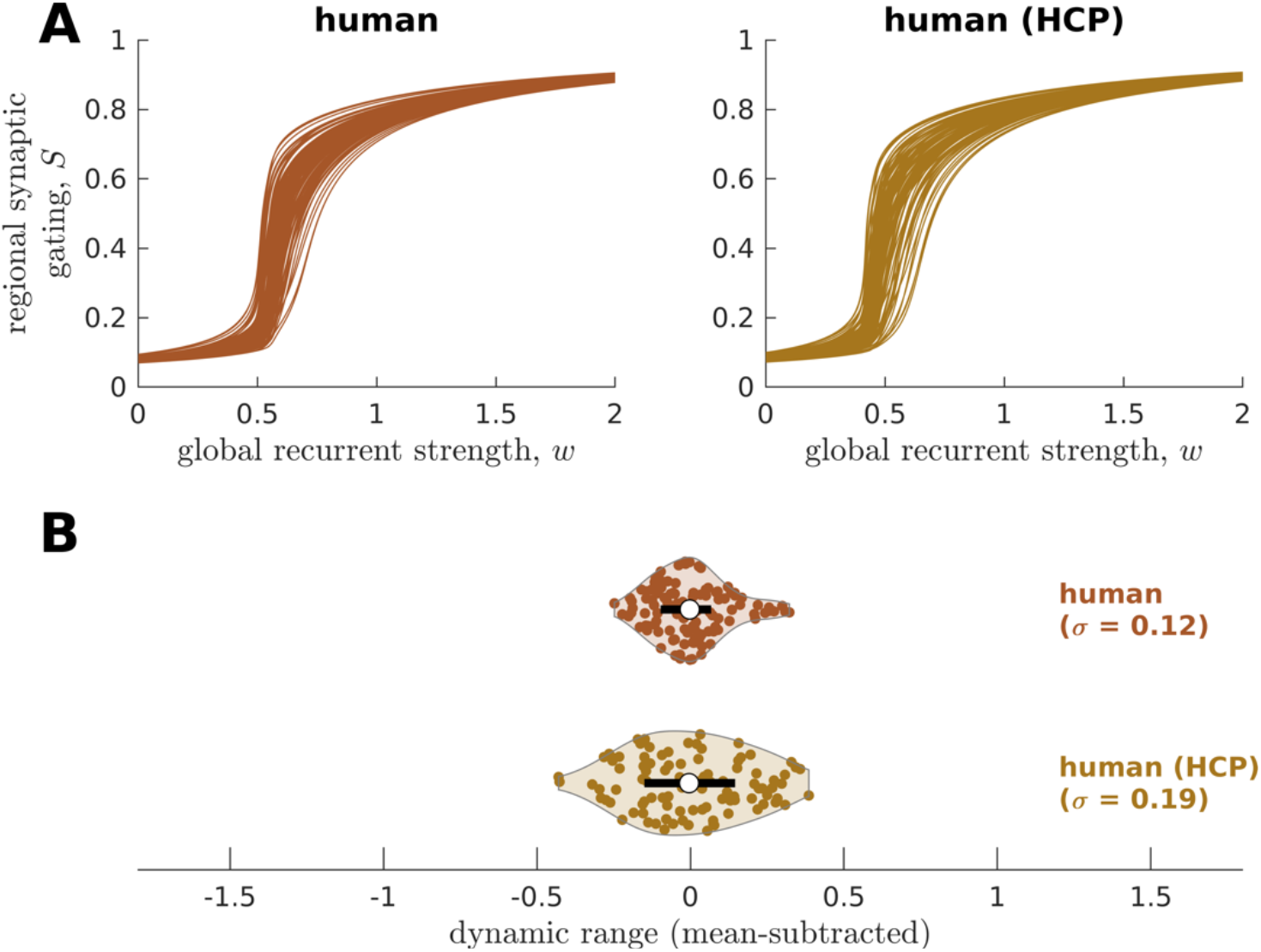
Replication of human neural dynamics on an independent dataset. (**A**) Regional neural dynamics as a function of global recurrent strength (*w*) for the original human connectome and human connectome obtained from the Human Connectome Project (HCP). (**B**) Violin plot of the distribution of dynamic ranges across brain regions. Each violin shows the first to third quartile range (black line), median (white circle), raw data (dots), and kernel density estimate (outline). The data are mean-subtracted for visual purposes. *σ* is the standard deviation of the distribution.

**Fig. S9.**
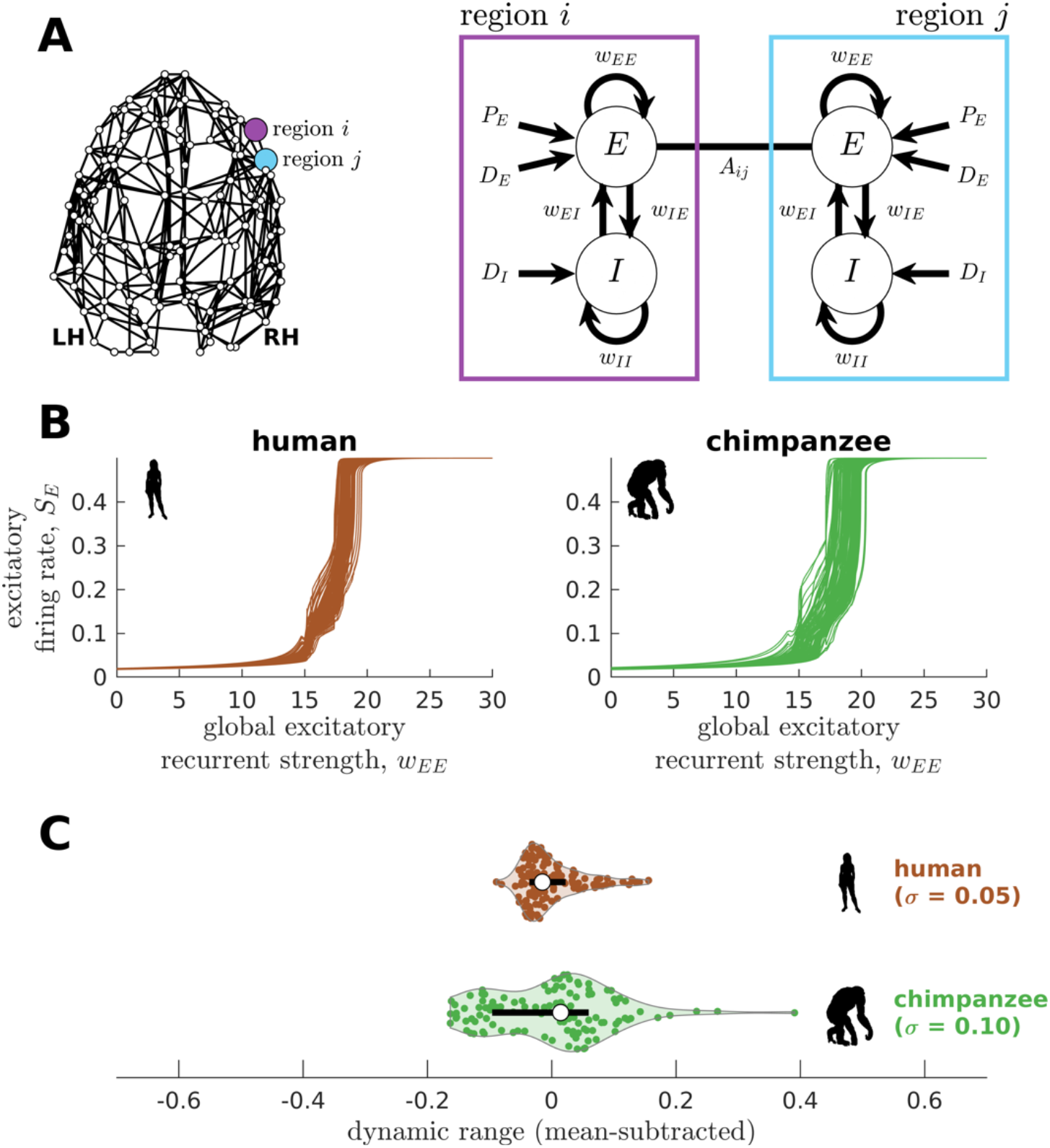
Replication of human and chimpanzee neural dynamics using a different biophysical model (the Wilson-Cowan model). (**A**) Exemplar connectome and schematic diagram of the Wilson-Cowan model. In this biophysical model, each brain region comprises interacting populations of excitatory (*E*) and inhibitory (*I*) neurons. Connections within and between populations are represented by the *w* parameters; e.g., *w_EE_* represents the excitatory recurrent connection strength. The excitatory neural population is driven by a constant excitatory input *P_E_* and white noise with standard deviation *D_E_*, while the inhibitory population is only driven by white noise with standard deviation *D_E_*. Regions *i*. and *j* are connected with weight *A_ij_* based on the connectomic data. (**B**) Regional neural dynamics (excitatory firing rate *S_E_*) as a function of global excitatory recurrent strength (*w_EE_*). (**C**) Violin plot of the distribution of dynamic ranges across brain regions. Each violin shows the first to third quartile range (black line), median (white circle), raw data (dots), and kernel density estimate (outline). The data are mean-subtracted for visual purposes. *σ* is the standard deviation of the distribution.

**Fig. S10.**
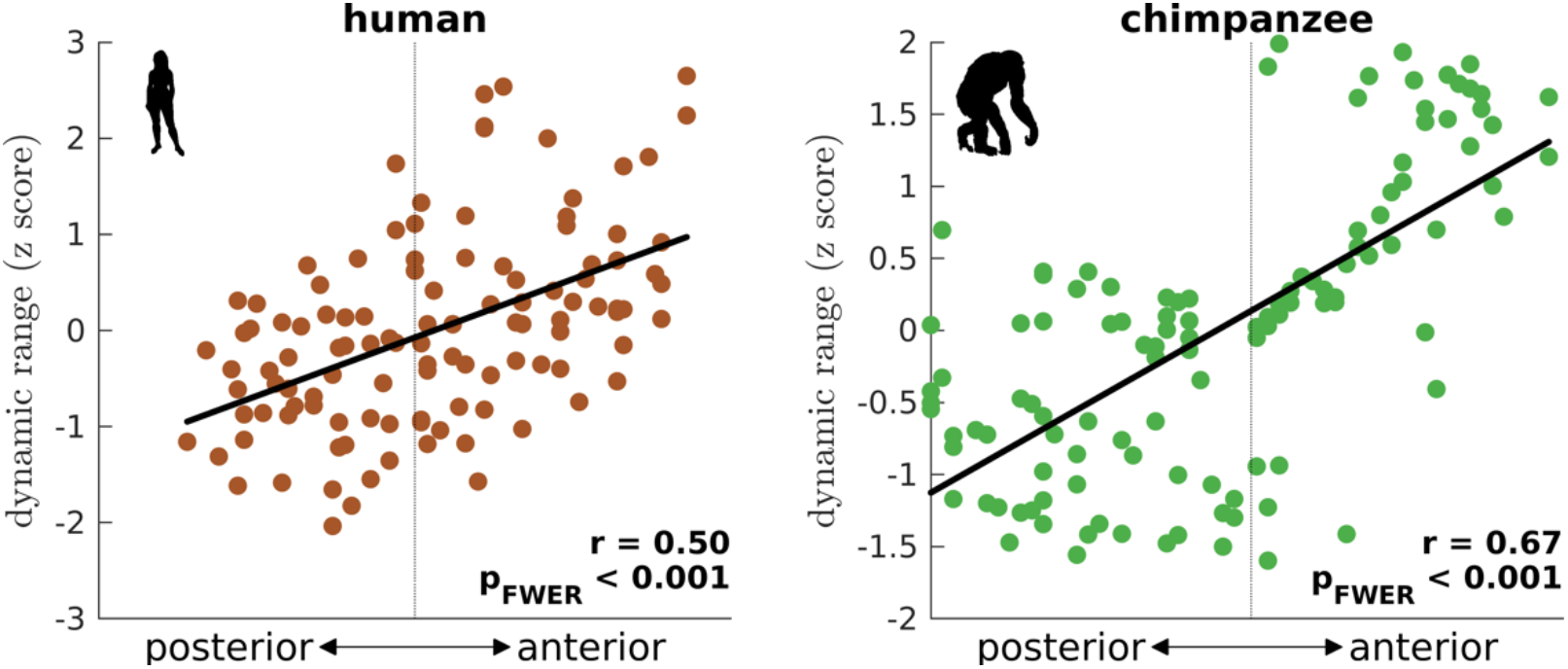
Gradient of dynamic ranges along the anterior-posterior axis. The dynamic range values are transformed to z scores. The solid line represents a linear fit with correlation coefficient r and FWER corrected p-value p_FWER_ after multiple comparisons.

**Fig. S11.**
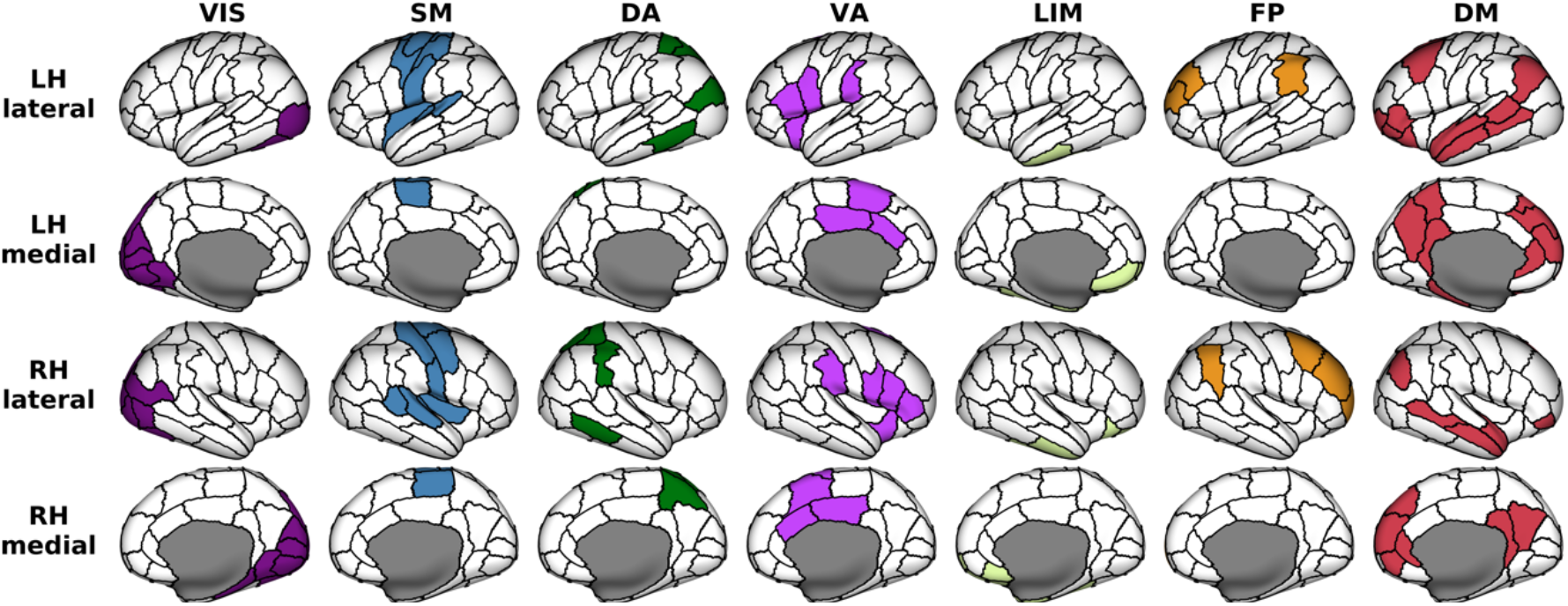
Anatomical locations of regions clustered according to seven canonical brain networks. VIS = Visual; SM = Somatomotor; DA = Dorsal Attention; VA = Ventral Attention; LIM = Limbic; FP = Frontoparietal; DM = Default Mode. These functional networks are mapped onto the 114-region atlas in Table S1. The networks are visualized on inflated human cortical surfaces.

**Fig. S12.**
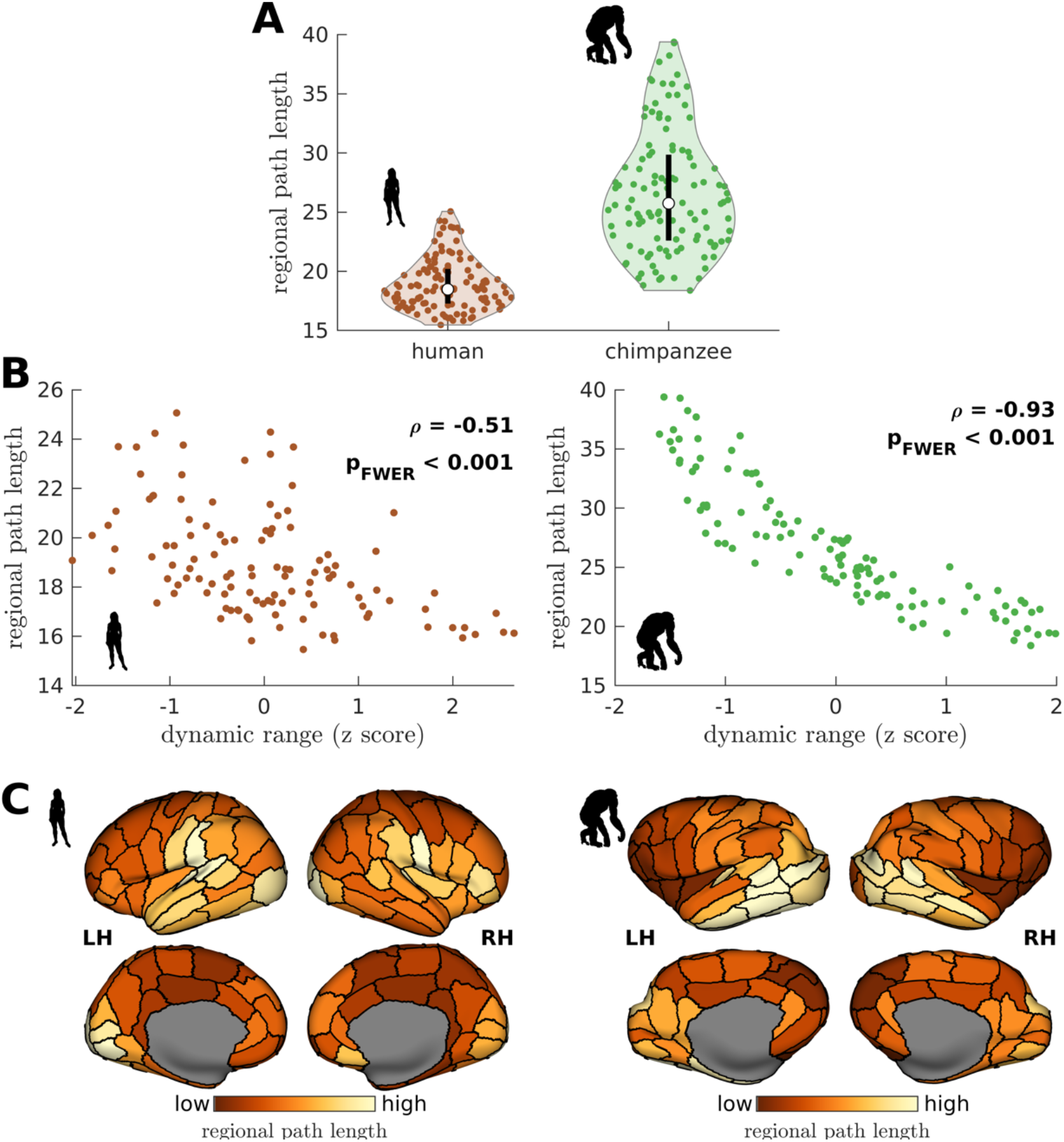
Human and chimpanzee connectome topological properties. (**A**) Violin plot of the distribution of regional path length across brain regions. Regional path length represents the average of the path lengths between a brain region and all other regions. Path length corresponds to the total topological distance of the shortest path between regions taking into account the weight of connections; topological distance = 1/weight. Each violin shows the first to third quartile range (black line), median (white circle), raw data (dots), and kernel density estimate (outline). (**B**) Average path length as a function of z-score-transformed dynamic ranges. *ρ* is the Spearman rank correlation and p is the FWER corrected p-value after multiple comparisons. (**C**) Spatial organization of regional path lengths. Data are visualized on inflated cortical surfaces (as per Fig. 2C). Light color represents high regional path length and dark color represents low regional path length.

**Fig. S13.**
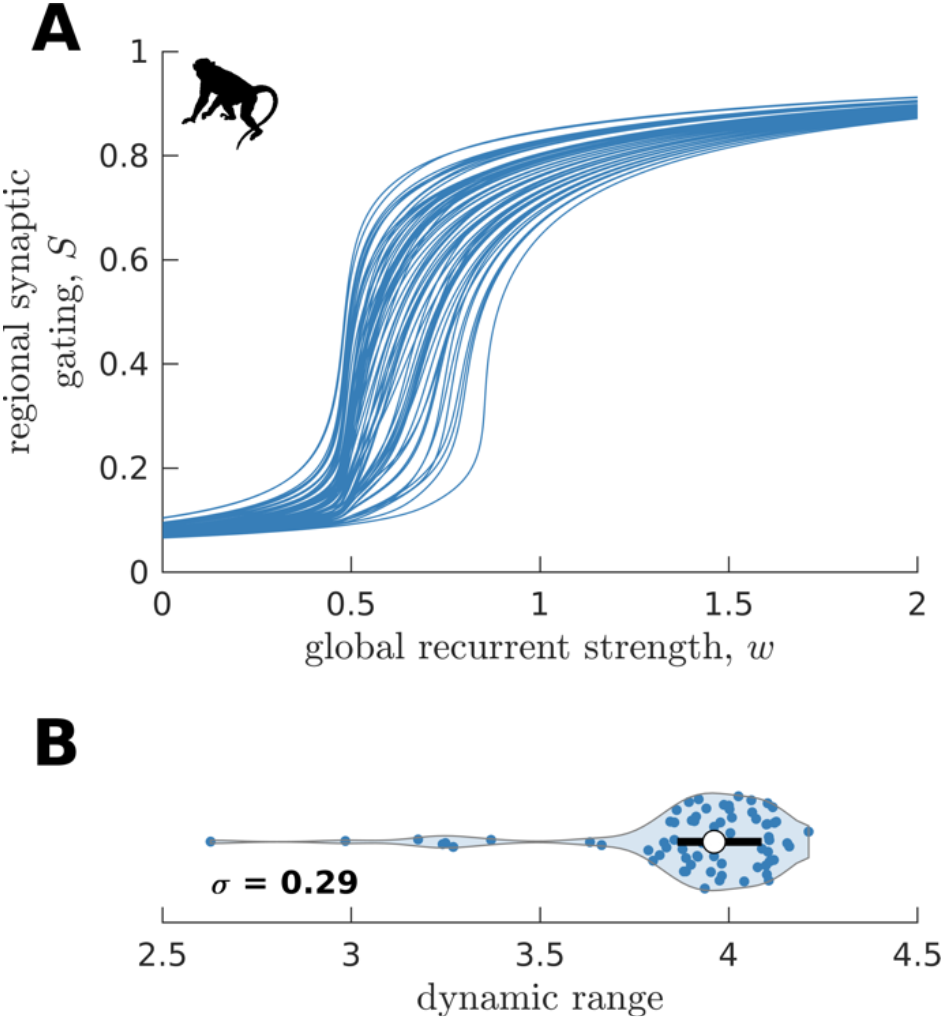
Replication of macaque neural dynamics on an independent dataset (CoCoMac). (**A**) Regional neural dynamics as a function of global recurrent strength (*w*). (**B**) Violin plot of the distribution of dynamic ranges across brain regions. The violin shows the first to third quartile range (black line), median (white circle), raw data (dots), and kernel density estimate (outline). *σ* is the standard deviation of the distribution.

**Fig. S14.**
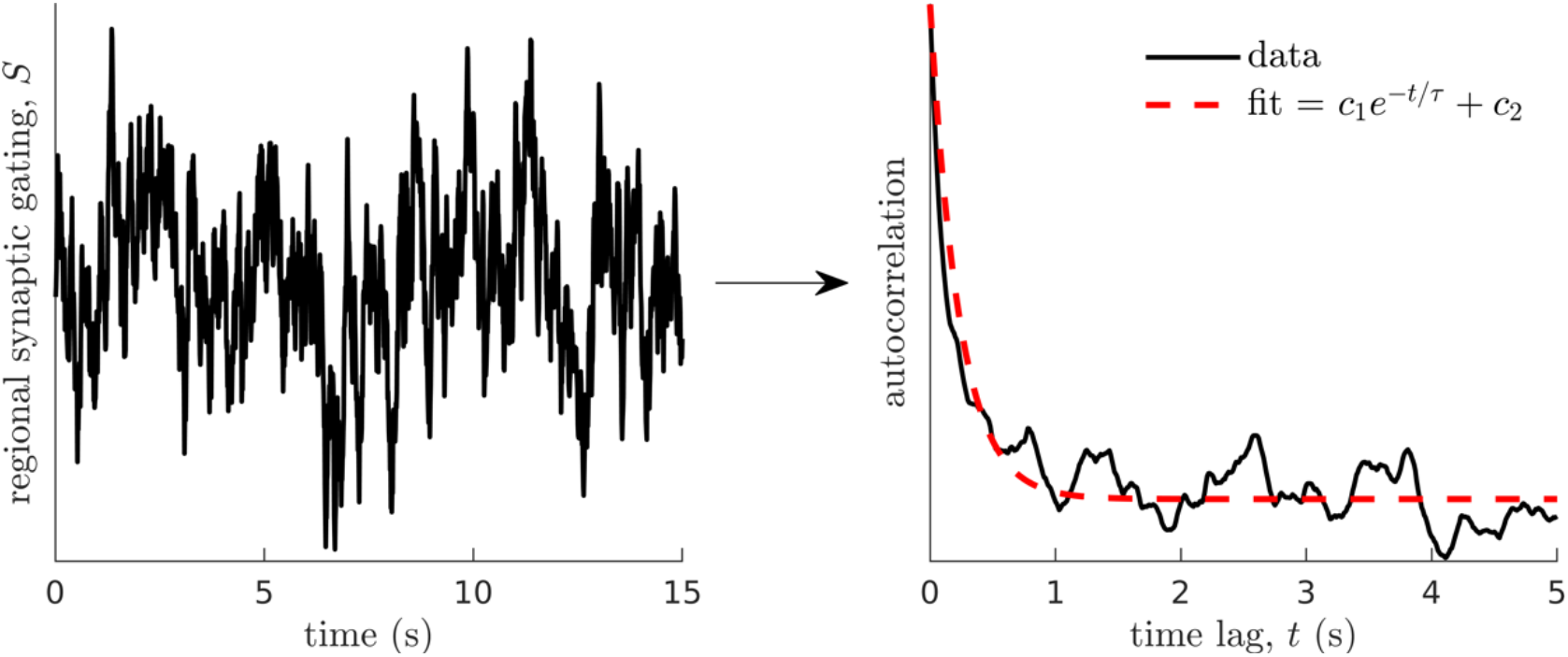
Method for calculating neural timescales. (Left) Sample regional neural activity. (Right) Autocorrelation of the data (neural activity) as a function of time lag (solid line) and corresponding exponential fit (dashed line) from which the timescale *τ* is estimated. Note that *c*_1_ and *c*_2_ are fitting constants.

**Fig. S15.**
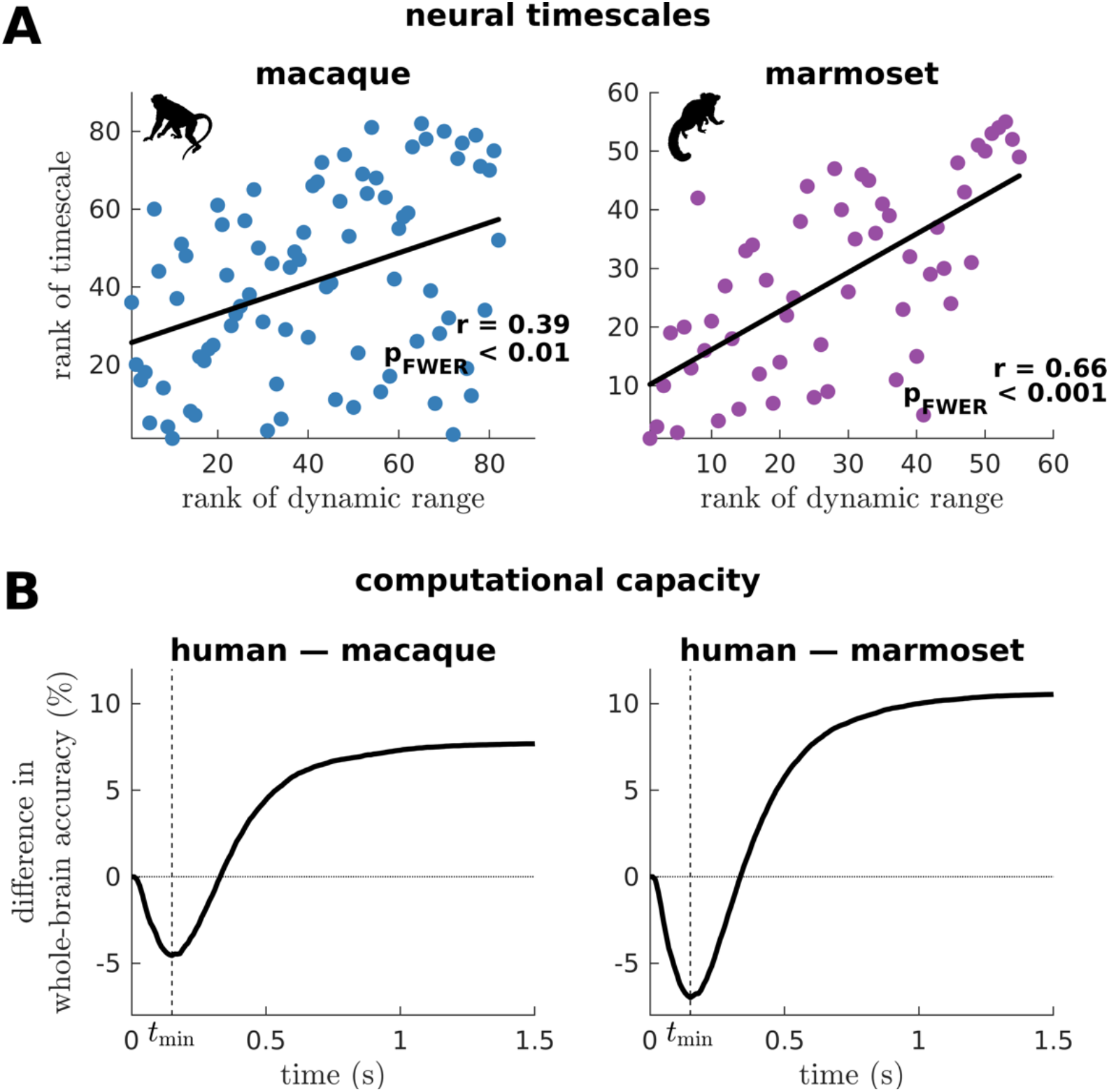
Macaque and marmoset neural timescales and their connectome’s computational capacity. (**A**) Ranked neural timescales as a function of ranked dynamic ranges (similar to Fig. 4A). The solid line represents a linear fit with correlation coefficient r and FWER corrected p-value p_FWER_ after multiple comparisons. (**B**) Human-macaque and human-marmoset difference in whole-brain accuracy across time (similar to Fig. 4E). The dashed line shows the time (*t*_min_) at which the difference in accuracy is most negative (i.e., macaque accuracy > human accuracy and marmoset accuracy > human accuracy).

**Fig. S16.**
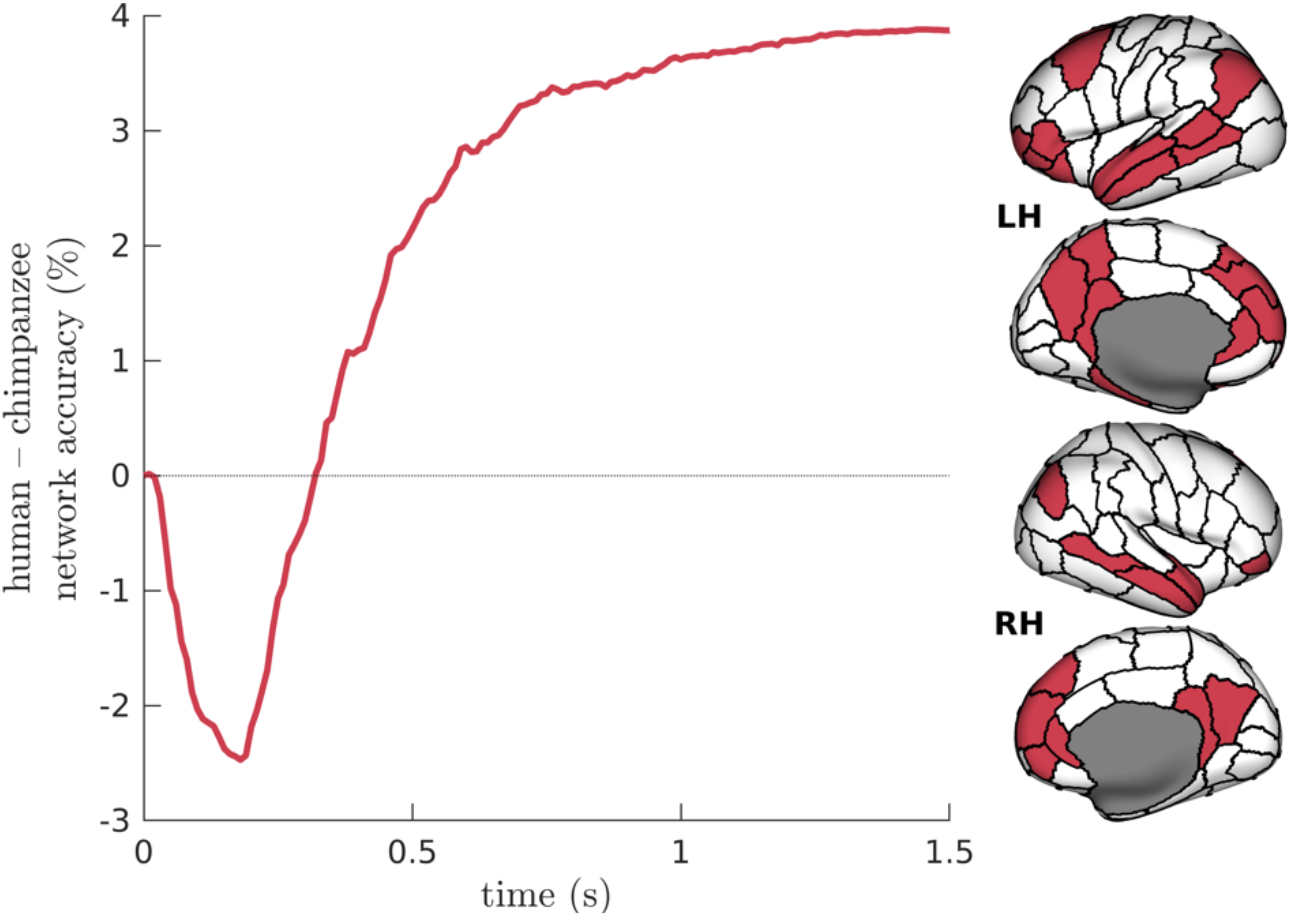
Difference in Default-Mode Network (DMN) accuracy across time between humans and chimpanzees. The DMN regions are visualized on inflated human cortical surfaces.

**Table S1.** List of 57 cortical regions in each hemisphere.

|  |  |  |
| --- | --- | --- |
| bankssts_1 | middle temporal_1 | rostral middle frontal_1 |
| caudal anterior cingulate_1 | middle temporal_2 | rostral middle frontal_2 |
| caudal middle frontal_1 | parahippocampal_1 | rostral middle frontal_3 |
| cuneus_1 | paracentral_1 | superior frontal_1 |
| entorhinal_1 | parsopercularis_1 | superior frontal_2 |
| fusiform_1 | parsorbitalis_1 | superior frontal_3 |
| fusiform_2 | parstriangularis_1 | superior frontal_4 |
| inferior parietal_1 | pericalcarine_1 | superior parietal_1 |
| inferior parietal_2 | postcentral_1 | superior parietal_2 |
| inferior temporal_1 | postcentral_2 | superior parietal_3 |
| inferior temporal_2 | postcentral_3 | superior temporal_1 |
| isthmus cingulate_1 | posterior cingulate_1 | superior temporal_2 |
| lateral occipital_1 | precentral_1 | supramarginal_1 |
| lateral occipital_2 | precentral_2 | supramarginal_2 |
| lateral orbitofrontal_1 | precentral_3 | frontal pole_1 |
| lateral orbitofrontal_2 | precentral_4 | temporal pole_1 |
| lingual_1 | precuneus_1 | transverse temporal_1 |
| lingual_2 | precuneus_2 | insula_1 |
| medial orbitofrontal_1 | rostral anterior cingulate_1 | insula_2 |

## Notes

### Competing Interest Statement

The authors have declared no competing interest.

